# Structure elucidation, biosynthetic gene cluster distribution, and biological activities of ketomemicin analogs in *Salinispora*

**DOI:** 10.1101/2024.10.29.620863

**Authors:** Gabriel Castro-Falcón, Dulce G. Guillén-Matus, Elany Barbosa Da Silva, Wentao Guo, Alicia Ross, Mateus Sá Magalhães Serafim, Thais Helena Maciel Fernandes, Dean J. Tantillo, Anthony J. O’Donoghue, Paul R. Jensen

## Abstract

We report three new ketomemicin pseudopeptides (**1**–**3**) from extracts of the marine actinomycete *Salinispora pacifica* strain CNY-498. Their constitution and relative configuration were elucidated using NMR, mass spectrometry, and quantum chemical calculations. Using GNPS molecular networking and publicly available *Salinispora* LCMS datasets, five additional ketomemicin analogs (**4**–**8**) were identified with ketomemicin production detected broadly across *Salinispora* species. The ketomemicin biosynthetic gene cluster (*ktm*) is highly conserved in *Salinispora*, occurring in 79 of 118 public genome sequences including eight of the nine named species. Outside *Salinispora, ktm* homologs were detected in various genera of the phylum Actinomycetota that might encode novel ketomemicin analogs. Ketomemicins **1**–**3** were tested against a panel of eleven proteases, with **2** displaying moderate inhibitory activity. This study describes the first report of ketomemicin production by *Salinispora* cultures, the distribution of the corresponding biosynthetic gene cluster, and the protease inhibitory activity of new ketomemicin derivatives.

**Table of Content/Graphical Abstract:** 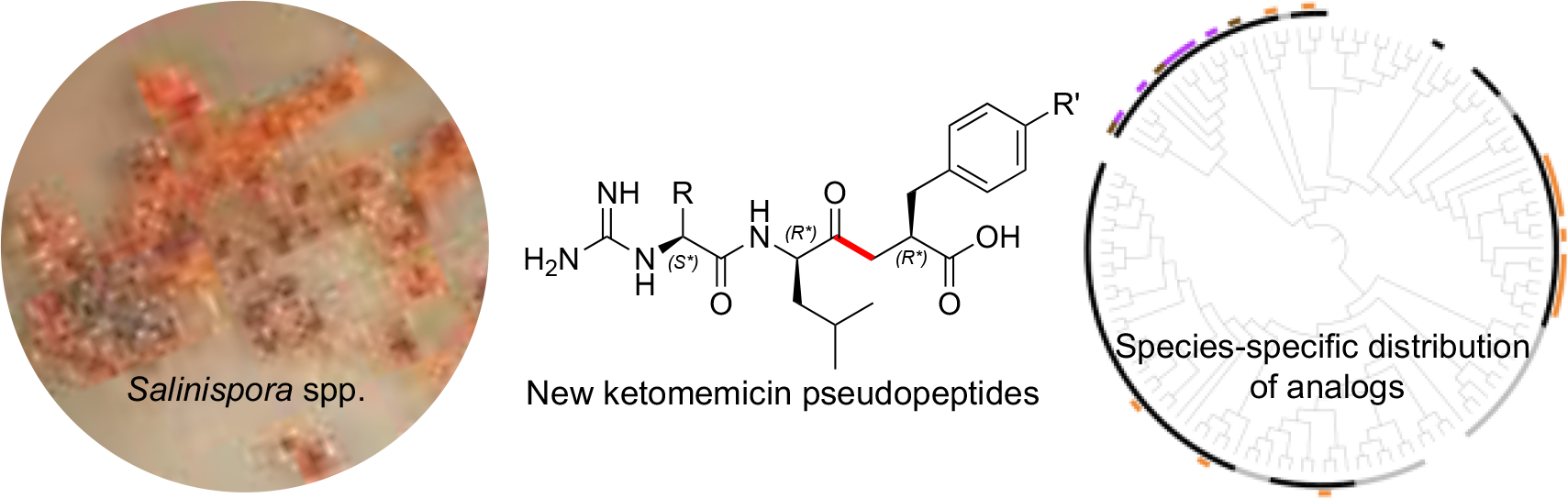

## INTRODUCTION

The pseudopeptide natural products ketomemicin A, B1–B6, and C were previously discovered following heterologous expression of biosynthetic gene clusters (BGCs) from *Micromonospora* sp. ATCC-39149, *Streptomyces mobaraensis* NBRC 13819, and *Salinispora tropica* CNB-440, respectively (**Figure 1**).^1^ The six-gene BGCs, named *ktm*, encode an aldolase (*ktmA*), a PLP-dependent amino acid C-acyltransefrase (*ktmB*), a dehydratase (*ktmC*), a peptide ligase (*ktmD*), an amidinotransferase (*ktmE*), and a dehydrogenase (*ktmF*), and are thus independent of the more traditional ribosomal and non-ribosomal mechanisms of peptide natural product biosynthesis.^1,2,3^Ketomemicins have not been previously reported from *Salinispora* strains^4^nor were they detected in culture extracts of *S. tropica* CNB-440,^1^suggesting the BGC remained silent under the laboratory growth conditions employed.

**Figure 1.**
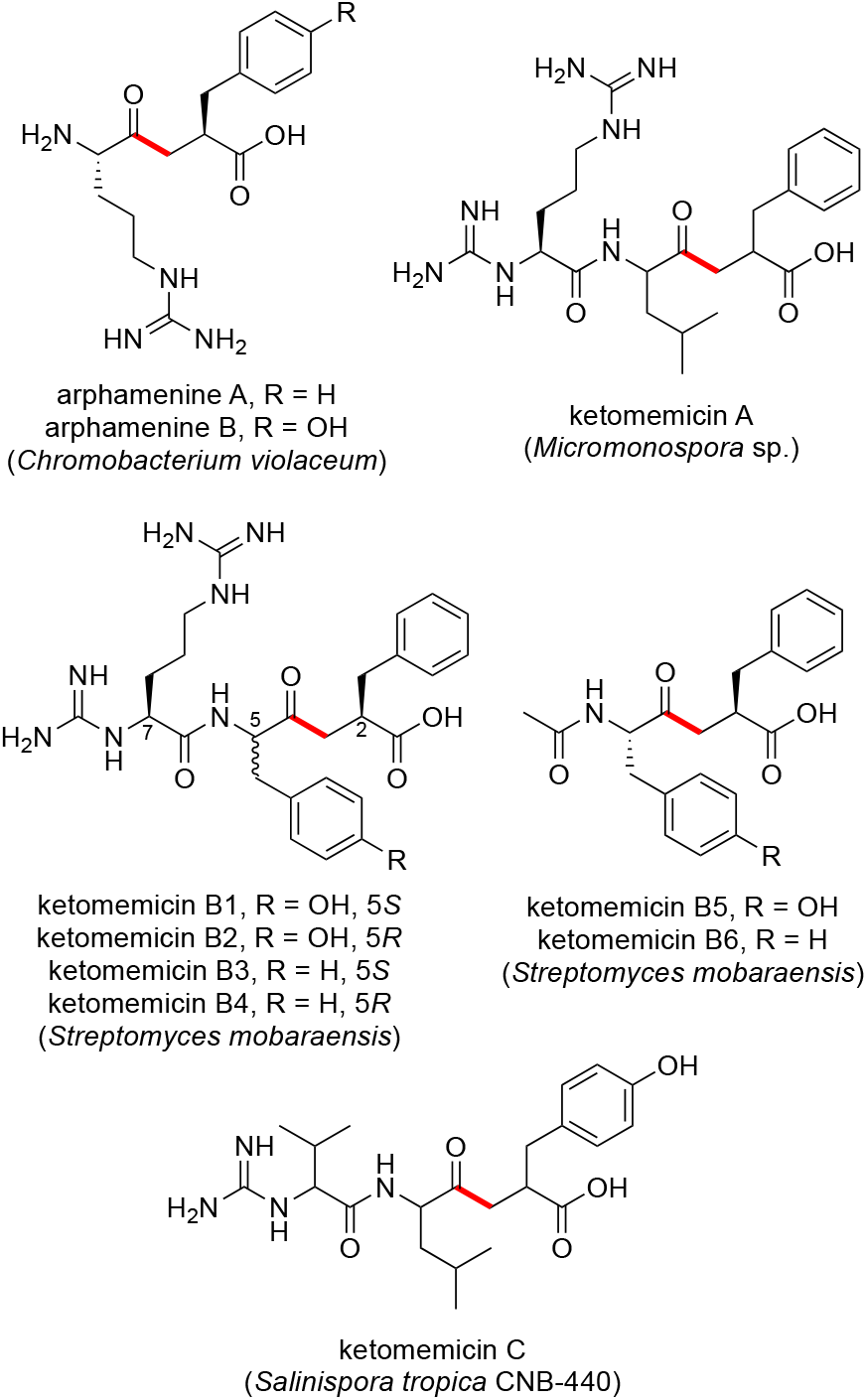
Previously reported ketomethylene-containing pseudopeptide natural products arphamenines and ketomemicins. The ketomethylene bond in each structure is shown in red. Producing organisms are shown in parentheses.

The natural products arphamenine A and B are structurally similar to the ketomemicins. They were discovered from the Gram-negative bacterium *Chromobacterium violaceum* due to their ability to inhibit the mammalian protease aminopeptidase B.^5,6^ Both ketomemicins and arphamenines contain amino acid residues typical of peptides but are considered pseudopeptides due to the presence of a ketomethylene bond in lieu of a typical peptide bond. Although ketomemicins and arphamenines are the only known naturally occurring ketomethylene-containing pseudopeptides, synthetic peptides with similar structures have been developed as protease inhibitors.^7^ Interestingly, the isosteric replacement of a peptide bond to a ketomethylene bond may be an evolved strategy of natural product protease inhibitors.^8^

However, we are unaware of any prior data describing the effects of the ketomemicins on protease activity.

In this work, we report the structures, relative configuration, and protease inhibitory activities of three novel ketomemicins (**1**–**3**) obtained from culture extracts of *Salinispora pacifica* CNY-498. We evaluated the production of ketomemicin analogs across *Salinispora* metabolomic datasets and assessed the diversity and distribution of the *ktm* BGC in the genus *Salinispora* and, more broadly, in the phylum Actinomycetota to show that additional diversity likely remains to be discovered within this unusual compound class.

## RESULTS AND DISCUSSION

### Isolation and Structure Elucidation of Ketomemicins

HPLC-MS screening of *Salinispora* culture extracts revealed three compounds produced by *S. pacifica* CNY-498 that were suggestive of a new series of natural products. To obtain enough of these compounds for NMR structure elucidation and biological testing, strain CNY-498 was grown in 18 x 1L cultures in A1FB medium with the addition of the adsorbent resin XAD-7 at day 8. The organic eluent from the collected resin and cells was subjected to C_18_ flash chromatography using a six-step solvent gradient of H_2_O and MeCN resulting in a fraction enriched in the three target compounds. This fraction was subjected to preparative HPLC to yield 1.0–0.4 mg of compounds **1**–**3**. Structure elucidation using HRMS and NMR spectroscopic analysis revealed that all three compounds were new derivatives of the natural product ketomemicin C, herein named according to their respective molecular mass as ketomemicin C-418 (**1**), ketomemicin C-432A (**2**), and ketomemicin C-432B (**3**) (**Fig. 2** and **Supplementary Figures S1-S15**).

**Figure 2.**
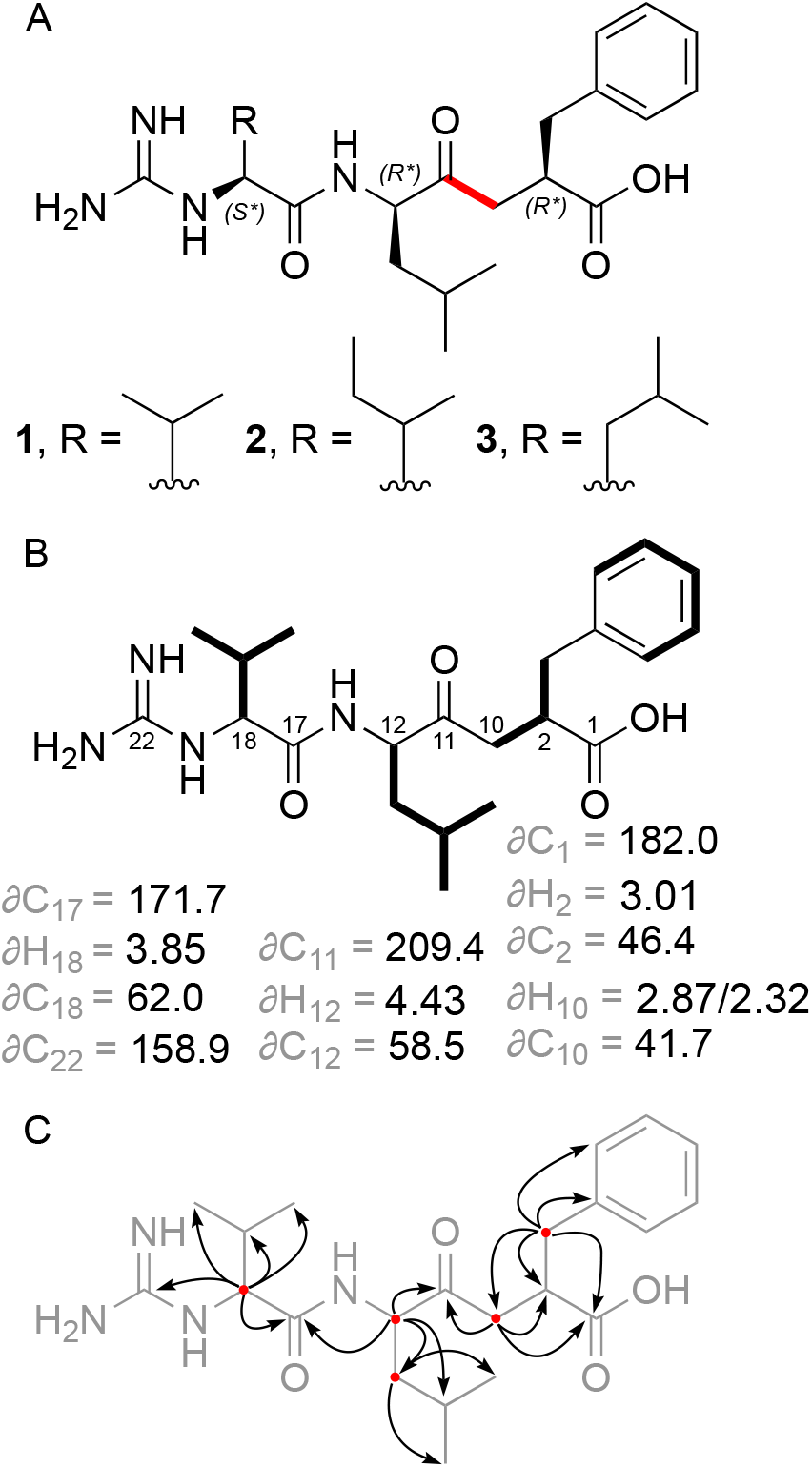
A) Structures of new ketomemicins (**1**–**3**) isolated in this work (ketomethylene bond in red). B) ^1^H and ^13^C NMR chemical shifts (in ppm) of backbone atoms and ^1^H–^1^H COSY correlations (bolded bonds) observed for 1.C) Key HMBC correlations observed for **1** represented as arrows from ^1^H to ^13^C atoms.

Ketomemicin C-418 (**1**), isolated as a thin white film, was analyzed by HRMS to give the molecular formula C_22_H_34_N_4_O_4_ (observed 419.2650 *m/z* [M+H]^+^, calculated 419.2653, -0.67 ppm error). In CD_3_OD, the ^1^H NMR spectrum indicated aromatic protons (δH 7.15, 7.22, 7.24 ppm), alpha-protons (δH 2.88, 2.32 ppm), deshielded aliphatic protons (δH 2.30/2.88, 2.99, 2.64/3.05 ppm), and shielded aliphatic protons (δH 0.91– 2.18 ppm), totaling 28 hydrocarbon protons. The remaining six protons exchanged with the deuterated NMR solvent and could not be detected. Notably, one of the ketomethylene protons (δ 2.88 ppm) showed a relatively diminished peak area due to partial exchanged with deuterium. HSQC and HMBC spectra revealed all 22 carbons in **1**, including a ketone (δC 209.4 ppm), carboxylic acid (δC 182.0 ppm), amide (δC 171.7 ppm), carbonyl alpha-carbons (δC 41.7, 46.4, 62.0 ppm), aromatics (δC, 126.9–141.5 ppm), guanidine (δC 158.9 ppm), and aliphatic carbons (δC 17.8–39.7 ppm). COSY and HMBC spectra respectively showed four spin systems and their interconnectedness (**Fig. 2A-C**). Compound **1** resembles the tripeptide Val-Leu-Phe but with a ketomethylene replacement (C_10_ – C_11_) and a guanidine group at the N-terminal valine.

Ketomemicin C-432A (**2**) and ketomemicin C-432B (**3**) were also isolated as thin white films. Their HRMS analysis indicated the molecular formula C_23_H_36_N_4_O_4_(calculated 433.2810 for M+H^+^) due to observed values of 433.2820 and 433.2824 *m/z* for the respective isomers. The NMR spectra for both **2** and **3** closely resembled that of **1**, except for signals related to the N-terminal amino acid, which indicated the presence of an isoleucine in **2** and a leucine in **3**. While the MS^2^spectra of **2** and **3** were very similar, the spectrum of **2** exclusively displayed a small fragment ion at 69.1 m/z indicative of the isoleucine residue (**Fig. S5** and **S10**).^9^

The relative configuration of **1** was determined by comparing the experimental ^1^H and ^13^C NMR chemical shift values to those calculated for the four possible diastereomers of **1** (as the guanidinium-carboxylate zwitterion) using quantum chemical computations.^10^Distinguishing the correct diastereomer presented a substantial challenge due to the flexibility of the molecules and the quantity of polar groups they contain. For each diastereomer, conformational searching was conducted using xTB-CREST to identify low-energy conformers.^11^These low-energy conformers were then optimized using *Gaussian16* at the restricted B3LYP-D3(0)/6-31+G(d,p) level of theory with an implicit solvation model (IEFPCM).^12-16^NMR chemical shift calculations were then performed for the lowest energy conformers within a 3 kcal/mol energy window using mPW1PW91/6-311+G(2d,p), with methanol as solvent.^17-18^The isotropic shielding values obtained from these calculations were converted to chemical shifts using scaling factors from the CHESHIRE dataset.^10^ The computed chemical shifts of the conformers for each diastereomer were weighted and averaged based on their relative free energies at the IEFPCM(methanol)-B3LYP/6-31+G(d,p) level using a script provided by Hoye and co-workers.^19^A comparison was made between the experimentally determined and the predicted chemical shifts of the candidate diastereomers. However, due to the similarity of the predicted chemical shifts for the four diastereomers, the conventional criteria of root-mean-square deviation (RMSD) and mean absolute error (MAE) were unable to provide a definitive assignment; all candidates exhibited a strong correlation between the experimental and computation NMR data, having only small deviations and no large outliers. Thus, a DP4+ analysis was conducted to obtain a more robust confidence analysis for the four diastereomers.^20^This analysis revealed that the (2R^*^, 12R^*^, 18S^*^) diastereomer was the best match to the experimental chemical shifts, with a computed probability of >93% when considering both ^1^H and ^13^C signals. Therefore, we consider this relative configuration to be the most probable. Additional details can be found in the Supporting Information (**Tables S4-S7**). The same relative configuration was assumed for **2** and **3** due to the almost identical NMR chemical shifts and specific optical rotation values of **1**–**3**.

### Diversity and Distribution of Ketomemicins in *Salinispora*

Using GNPS molecular networking,^21^ we queried for ketomemicin analogs with similar MS^2^ spectra to **1**–**3** in published LC-MS/MS datasets from *Salinispora* spp.^22-24^ This led to the identification of five ketomemicin analogs (**4**–**8**) in the Crüsemann et al. (2017) dataset, which includes organic extracts of 118 genome-sequenced *Salinispora* strains grown on agar (**Fig. S16**). The constitution of **4**–**8** could be putatively assigned by comparing their MS^2^ spectra with that of **1**–**3** (**Fig. 3**). While **6** is identical in constitution to the previously reported ketomemicin C (herein referred to as ketomemicin C-434)^1^ and **7** was previously reported based on the analysis of MS data,^2^ **4, 5**, and **8** are new compounds. Notably, the ketomethylene bond in all arphamenines and ketomemicins discovered to date is associated with a C-terminal phenylalanine-or tyrosine-derived residue.

**Figure 3.**
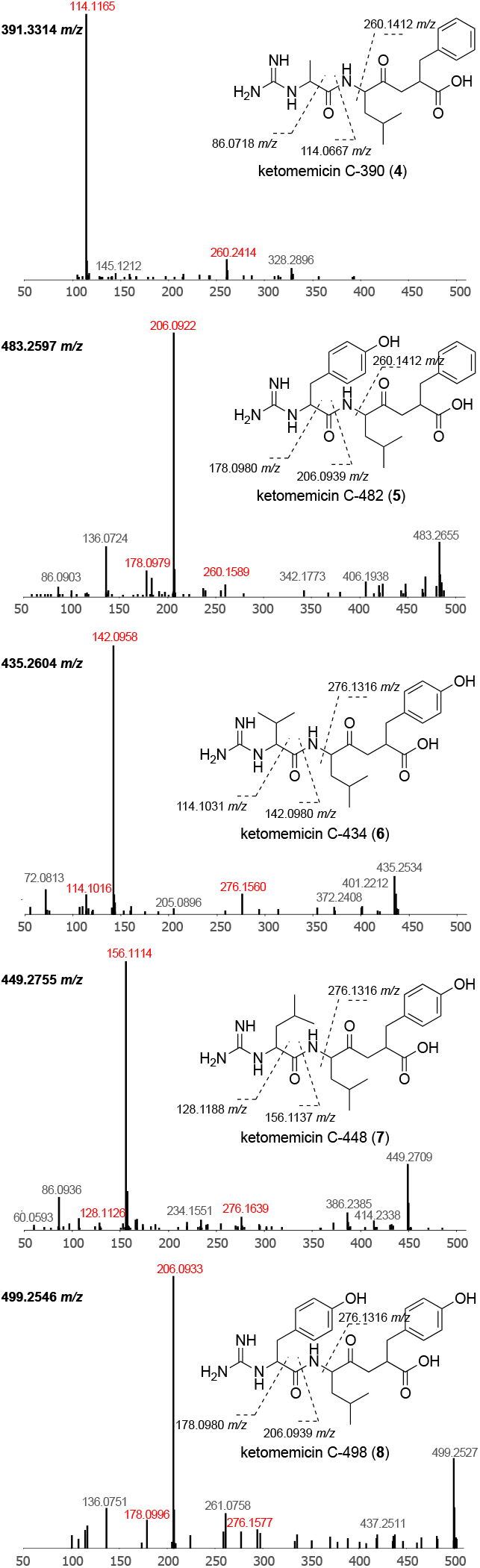
Structures and MS^2^ spectra of ketomemicins **4**–8.Characteristic mass fragments (in red) putatively arise from the cleavage of bonds crossed with dashed lines.

We next analyzed for ketomemicins across the Crüsemann et al. (2017) dataset using the “targeted feature detection” function within MZmine and the mass, retention time, and fragment ions as defining features for each metabolite.^25^From this analysis, we observed the production of **1**–**8** in 25 of 118 *Salinispora* strains (**Table S4**), corresponding to six of the nine currently described *Salinispora* species.^26^ The vast majority of these strains (19/25) were *S. tropica* and *S. pacifica*. When mapped on a maximum-likelihood phylogeny generated using 2,011 core genes from 118 *Salinispora* genomes,^26^ ketomemicin production was widely observed in *S. tropica* and more localized to specific clades within *S. pacifica* (**Fig. 4A**). Furthermore, species-specific production patterns were observed as ketomemicins **6**–**8** with the C-terminal tyrosine-derived residue were mainly produced by *S. tropica* while ketomemicins **1**–**4** with the C-terminal phenylalanine-derived residue were mainly produced by the other *Salinispora spp*., in particular *S. pacifica* (**Fig. 4A** and **Table S4**). Notably, only one of three *S. mooreana* strains produced ketomemicins and it yielded the highest levels of **1**–**3** across the entire dataset, while *S. arenicola* and *S. oceanensis* showed low and infrequent production of **1**–**8** (observed in 3/61 and 1/13 strains, respectively). Compound **5** was only seen in one of two *S. fenicalii* strains while ketomemicin production (**1**–**8**) was not observed in *S. cortesiana, S. goodfellowii*, or *S. vitiensis*. Together, these analyses revealed the broad yet inconsistent production of ketomemicins across the genus *Salinispora*. We speculate that ketomemicins have previously eluded detection due to their relatively low production levels.

**Figure 4.**
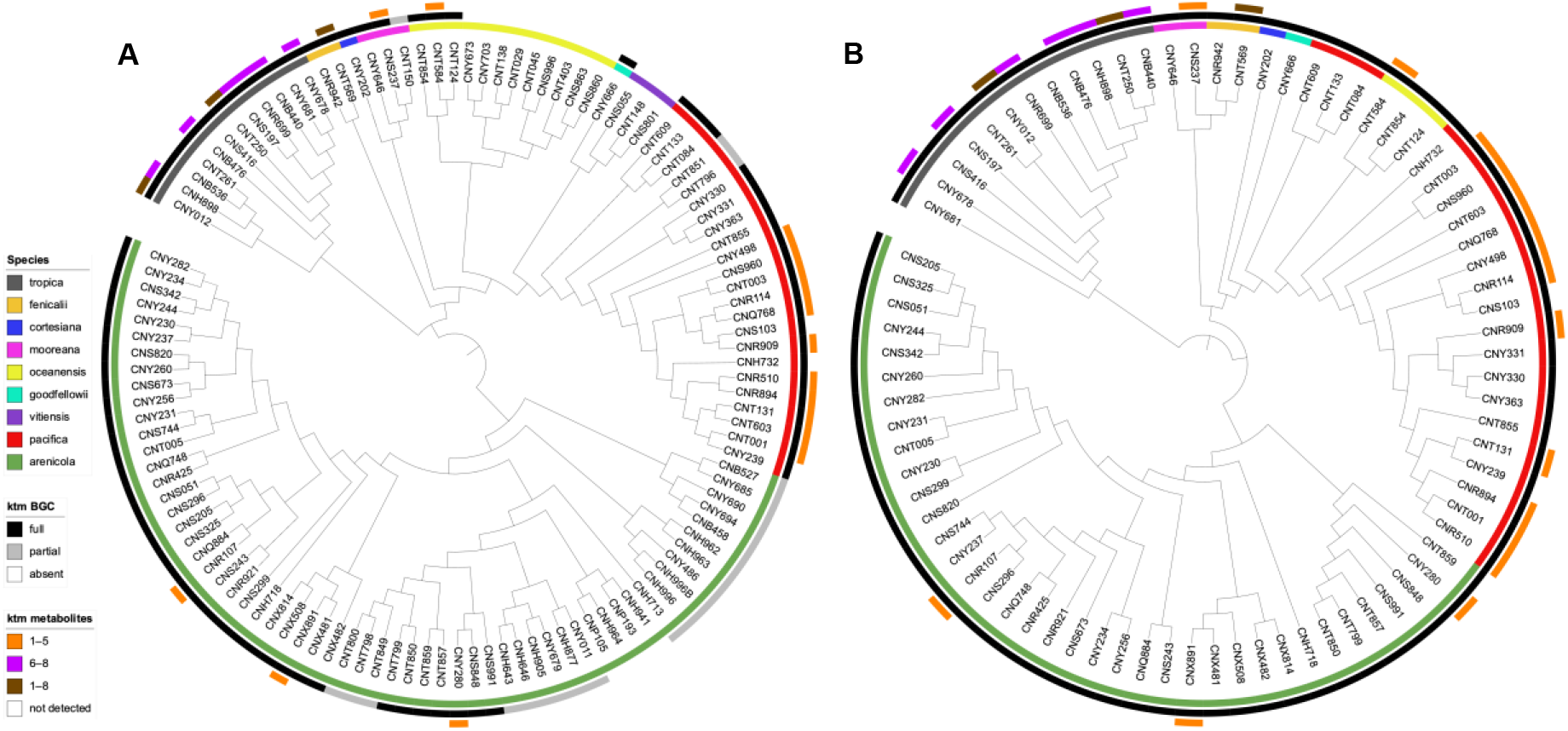
Phylogenetic relationships of the *ktm* BGC and ketomemicin production in *Salinispora*. A) Phylogenomic tree of 118 *Salinispora* strains representing all nine currently described species (inner circle), distribution of the *ktm* BGC (middle circle), and observed production of ketomemicins with C-terminal phenylalanine-derived residue (**1**– **5**), C-terminal tyrosine-derived residue (**6**–**8**) or both (**1**–**8**) (outer circle). B) Phylogeny (bootstrap value of 1000) of the complete *ktm* BGC observed in 79 *Salinispora* strains.

### Diversity and Distribution of *ktm* in *Salinispora* spp. and Actinomycetota

Using antiSMASH,^27^we detected high percent similarity homologs of all six ketomemicin biosynthetic genes (*ktmA*-*F*) reported from *Streptomyces mobaraensis* NBRC 13819^1^ in *S. pacifica* CNY-498. Using the *S. pacifica* BGC as input, we queried 118 *Salinispora* genomes using Cblaster^28^ and identified all six *ktm* genes (≥87% identity and 97% coverage) in 79 *Salinispora* strains spanning eight of the nine species (**Fig. 4A** and **Fig. S17)**. Interestingly, we also detected incomplete or partial *ktm* clusters containing two to five *ktm* genes (≥87% identity and 54% coverage) in 23 *Salinispora* genomes (20 *S. arenicola*, two *S. pacifica*, and one *S. mooreana*). These partial gene clusters were not on contig edges and thus do not appear to be sequencing artifacts. Similar observations of incomplete BGCs have been made for the desferrioxamine BGC (*des*) in *Salinispora*.^29^ Using Clinker,^30^ we observed high gene synteny among the *ktm* BGCs, although species-specific differences in the flanking genes suggests they may occur in different genomic environments (**Fig. 5** and **Fig. S17**), as reported for other *Salinispora* BGCs.^31^ A *ktm* BGC phylogeny generated using all six genes from the 79 *Salinispora* genomes was highly congruent with the phylogenomic tree (**Fig. 4B**), suggesting that *ktm* was present in the *Salinispora* common ancestor and has largely been passed down through vertical transmission. One exception is observed for *S. oceanensis* strains, which appear to have acquired the BGC from *S. pacifica* based on their position within the *S. pacifica* clade. When examining the relationships between the *ktm* BGC and ketomemicin production (**Fig. 4A**), compounds were only detected in 25 (31.6%) of the 79 strains with the six gene operon. In *S. arenicola*, they were only detected in 3 (0.08%) of 37 strains. It remains unclear if the BGCs that could not be linked to compound production are non-functional or are under different regulatory control. There was no evidence of the former based on comparative sequence analysis. As expected, ketomemicins were not detected in any of the strains with a partial *ktm* BGC.

**Figure 5.**
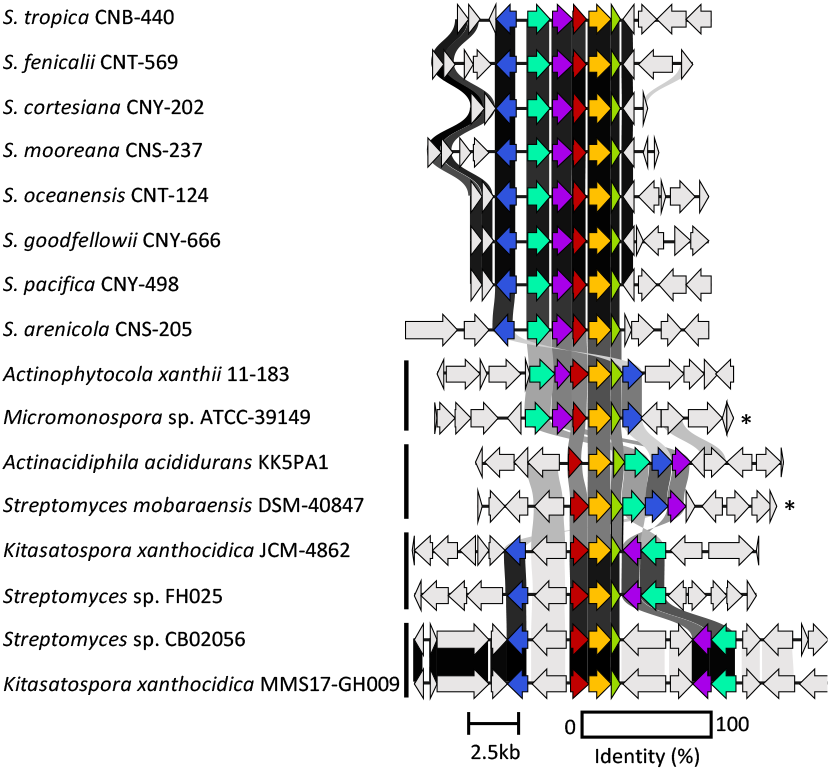
Synteny plot showing *ktm* and *ktm-*like biosynthetic gene clusters in *Salinispora* and diverse *Actinomycetota*. Representative *ktm* BGCs from eight *Salinispora* spp. are highly conserved across the genus (see **Figure S17** for a full list). Four additional versions of the BGC (vertical bars) were observed among 28 *Actinomycetota* strains (see **Figure S18** for a full list). Genes are colored as: *ktmA* (red), *ktmB* (yellow), *ktmC* (olive), *ktmD* (cyan), *ktmE* (blue), and *ktmF* (purple). Asterisks (^*^) denote experimentally validated *ktm* clusters outside of *Salinispora*.

We next used Cblaster to further assess the diversity and distribution of the *S. arenicola* CNY-498 *ktm* BGC within the NCBI genome database. We identified 28 non-*Salinispora* Actinomycetota that contain a BGC with homologs of *ktmA-F* (**Figure S18**), including *Micromonospora* sp. ATCC-39149 and *Streptomyces mobaraensis* NBRC 13819 from which the *ktm* BGCs were heterologously expressed (**Figure 1**).^1^ These sequences could be grouped into four *ktm*-like BGC types based on gene synteny (**Figure 5**). While the products of two of these have been experimentally validated, the other two could yield new ketomemicin or ketomethylene-containing pseudopeptide natural products.

### Biological Activities of Ketomemicins

Due to their structural similarity to the arphamenines, which are known protease inhibitors,^5,6^ **1**–**3** were tested at 10 μM against a panel of eleven proteases of diverse origins, including humans (cathepsin B, D, L, aminopeptidase B, and human 20S proteasome), parasites (cruzain and *Trypanosoma brucei* cathepsin L), and viruses (SARS-CoV, SARS-CoV-2, and MERS-CoV main proteases, and papain-like protease) (**Table 1**). At these concentrations ketomemicins **1**– **3** were not active against aminopeptidase B, which is the target of the arphamenines. The only activity detected was for compound **2**, which displayed moderate inhibition against the main proteases (M^pro^) of SAR-CoV-1, SARS-Co-V-2, and MERS-Co-V as well as cruzain, while not being active against TbrCatL. Compounds **1**–**3** were also tested for antibacterial activity against *Escherichia coli* MG1655 and *Pseudomonas aeruginosa* and were inactive at the highest test concentration (32 μg/μL).

**Table 1.**
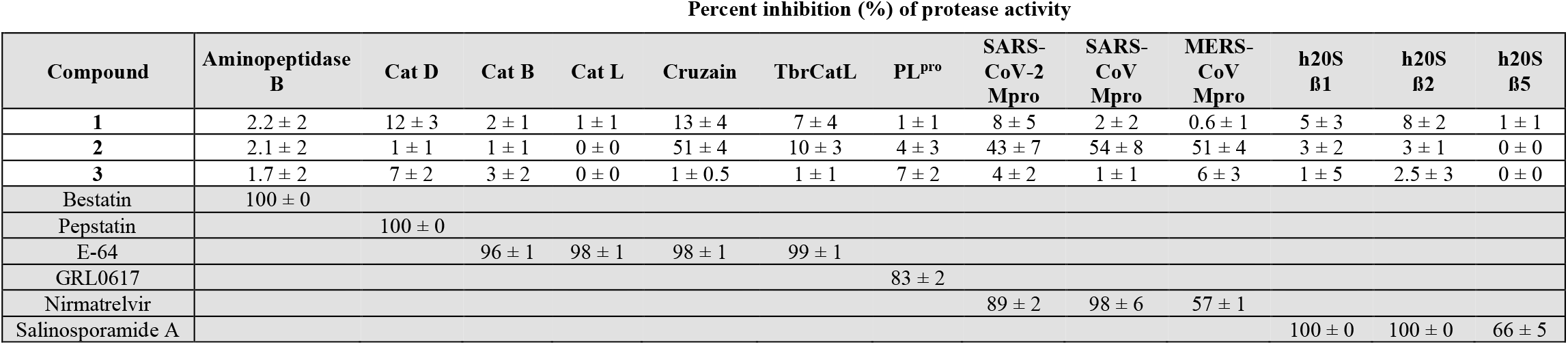
Inhibitory activities of ketomemicins 1–3 (at 10 μM) against a panel of eleven proteases. Average percent inhibition is reported for the mean of two independent experiments each performed in triplicate. Errors are given as the ratio of the standard deviation to the square root of the number of measurements. Control inhibitors (highlighted in grey) were tested at 10 μM, except for nirmatrelvir (tested at 100 nM). Cat B: Cathepsin B; Cat D: cathepsin D; Cat L: cathepsin L; TbrCatL: *Trypanosoma brucei* cathepsin-L like; PL^pro^: papain-like protease; h20S: human 20S proteasome.

In conclusion, we report the structures of new ketomemicins from cultures of the marine actinomycete *Salinispora pacifica* CNY-498. We describe the distribution of the ketomemicins and the ketomemicin BGC (*ktm*) in the paired metabolomic and genomic dataset of 118 *Salinispora* strains. We report two types of ketomemicin-like BGCs outside of *Salinispora* that might code for yet-to-be-characterized variants in this family of natural products. Finally, we report the inhibitory activities for the ketomemicins against a range of proteases.

## EXPERIMENTAL SECTION

### General Experimental Procedures

Optical rotations were recorded on a Jasco P-2000 polarimeter. UV spectra were measured on a Beckman-Coulter DU800 spectrophotometer. 1D and 2D NMR spectroscopic data were obtained on a JEOL 500 MHz or a Bruker 600 MHz NMR spectrometer. NMR chemical shifts were referenced to the residual solvent peaks (δH 3.31 and δC 49.15 for CD_3_OD). High-resolution ESI-TOF mass spectrometric data were acquired on an Agilent 6530 Accurate-Mass Q-TOF mass spectrometer coupled to an Agilent 1260 LC system.

### Cultivation

A frozen stock of *Salinispora pacifica* CNY-498 was inoculated into 50 mL of medium A1 [1% potato starch, 0.4% yeast extract, and 0.2% peptone in 2.2% InstantOcean®]. The seed culture was shaken at 200 rpm and 28*°*C for seven days then used to inoculate 1 L of medium A1 in a 2.8 L Fernbach flask. This culture was similarly shaken at 200 rpm and 28°C for eight days after which 20 mL were inoculated into each of 18 x 2.8 L Fernbach flasks containing 1 L of medium A1FB [A1 supplemented with 0.01% potassium bromide and 0.03% iron (III) sulfate (5·H_2_O)]. After eight days of shaking at 200 rpm and 28*°*C, 25 g of sterile XAD-7 adsorbent resin was added to each flask. After two additional days of cultivation, the 18 L were filtered through cheesecloth to collect the resin (and some cell material), which were soaked in acetone (3 L) for 2 h with gentle agitation. The acetone extract was filtered through a cotton plug and concentrated via rotary evaporation. The resulting solution was partitioned in a separatory funnel between EtOAc and H_2_O (1:1 mixture, 1 L total). The organic phase was collected, dried over anhydrous sodium sulfate, and concentrated via rotary evaporation to yield a red crude extract (500 mg).

### Isolation of ketomemicins

The organic extract was fractionated using C18 column flash chromatography (5g) and a six-step elution gradient from 100% H_2_O (0.1% formic acid) to 100% MeCN (0.1% formic acid) to yield six fractions. Fraction 4 (60% MeCN, 18.8 mg) was concentrated, resuspended, and separated over HPLC [mobile phase: 70% MeCN in H_2_O (0.1% formic acid) at 3 ml·min^-1^; stationary phase: 5 µm, C18(2), 100 Å, 250 x 10 mm (Phenomenex, Luna) column] to yield subfractions A (2-4 min, 3.8 mg) and B (4-10 min, 11.9 mg). Subfraction A was further separated by HPLC [mobile phase: 30% MeCN in H_2_O (0.1% formic acid) at 3 ml·min^-1^; stationary phase: 5 µm, C18(2), 100 Å, 250 x 10 mm (Phenomenex, Luna) column] to yield ketomemicin C-318 (**1**, t_R_ = 14 min, 0.8 mg), ketomemicin C.332A (**2**, t_R_ = 20 min, 0.7 mg) and ketomemicin C.332B (**3**, t_R_ = 22 min, 0.5 mg).

Ketomemicin C-418 (**1**): 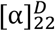; UV/vis (MeOH) λ (log ε) 200 (3.23), 212 (3.01) nm; ^1^H and 2D NMR, Table S1.

Ketomemicin C-432A (**2**): 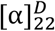; UV/vis (MeOH) λ (log ε) 200 (3.14), 212 (2.88) nm) nm; ^1^H and 2D NMR, Table S2.

Ketomemicin C-432B (**3**): 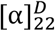; UV/vis (MeOH) λ (log ε) 200 (3.03), 212 (2.74) nm) nm; ^1^H and 2D NMR, Table S3.

## ASSOCIATED CONTENT

Additional experimental procedures, UV/vis, MS, and MS^2^ spectra, NMR spectroscopic data, additional details on quantum chemical computations, MS^2^ molecular networks, metabolite distribution analysis, and BGC synteny plots.

## AUTHOR INFORMATION

## Corresponding Author

Gabriel Castro-Falcón − Center for Marine Biotechnology and Biomedicine, Scripps Institution of Oceanography, University of California, San Diego, La Jolla, California 92093, United States

## Authors

Dulce G. Guillén-Matus − Center for Marine Biotechnology and Biomedicine, Scripps Institution of Oceanography, University of California, San Diego, La Jolla, California 92093, United States

Elany Barbosa Da Silva − Skaggs School of Pharmacy and Pharmaceutical Sciences, Center for Discovery and Innovation in Parasitic Diseases, University of California San Diego, La Jolla, California 92093, United States

Wentao Guo − Department of Chemistry, University of California Davis, Davis, California 95616, United States

Alicia Ross − Department of Chemistry, University of California Davis, Davis, California 95616, United States

Mateus Sá Magalhães Serafim − Skaggs School of Pharmacy and Pharmaceutical Sciences, Center for Discovery and Innovation in Parasitic Diseases, University of California San Diego, La Jolla, California 92093, United States

Thais Helena Maciel Fernandes − Skaggs School of Pharmacy and Pharmaceutical Sciences, Center for Discovery and Innovation in Parasitic Diseases, University of California San Diego, La Jolla, California 92093, United States

Dean J. Tantillo − Department of Chemistry, University of California Davis, Davis, California 95616, United States

Anthony J. O’Donoghue − Skaggs School of Pharmacy and Pharmaceutical Sciences, Center for Discovery and Innovation in Parasitic Diseases, University of California San Diego, La Jolla, California 92093, United States

Paul R. Jensen − Center for Marine Biotechnology and Biomedicine, Scripps Institution of Oceanography, University of California, San Diego, La Jolla, California 92093, United States; Phone: 858-534-7322;

## Notes

The authors declare no competing financial interest.

## ACNKOWLEDGMENTS

This work was supported by the National Institutes of Health (R01GM085770) to P.R.J. We thank the San Diego IRACDA postdoctoral program for funding to G.C.F. We thank B. Duggan from the UCSD SSPPS NMR Facility and A. Mrse from the UCSD Department of Chemistry and Biochemistry for assistance with NMR experiments, Y. Su from the UCSD Molecular Mass Spectrometry Facility for HRMS measurements, and P. Fajtova from UCSD Skaggs School of Pharmacy for assistance with the proteasome assay. We thank the CAPES Foundation (grant # 88887.595578/2020-00 and 88887.684031/2022-00) for funding to M.S.M.S. A high-resolution LC-MS instrument was provided by the National Institutes of Health (S10 OD0106400). Computational support from the NSF ACCESS program is gratefully acknowledge.

