## Supplemental Information for "Structure elucidation, biosynthetic gene cluster distribution, and biological activities of ketomemicin analogs in *Salinispora*"

**Table of Contents**

**General Experimental**……………………………………………………………………………………3

**Table S1.** NMR table for ketomemicin C-418 (**1**)…………………..……………….……………..………5

**Figure S1.** [1]H NMR spectrum of ketomemicin C-418 (**1**) (600 MHz, $CD_3OD$)………...…………………6

**Figure S2.** COSY spectrum of ketomemicin C-418 (**1**) (600 MHz, $CD_3OD$)…………...………………7

**Figure S3.** HSQC spectrum of ketomemicin C-418 (**1**) (600 MHz, $CD_3OD$)………...……………………8

**Figure S4.** HMBC spectrum of ketomemicin C-418 (**1**) (600 MHz, $CD_3OD$)…..…………………………9

**Figure S5.** HRMS and MS/MS spectra of ketomemicin C-418 (**1**)……………..………………………..10

**Table S2.** NMR table for ketomemicin C-432A (**2**)…………………………………..…………….....11

**Figure S6.** [1]H NMR spectrum of ketomemicin C-432A (**2**) (600 MHz, $CD_3OD$).……………………..…12

**Figure S7.** COSY spectrum of ketomemicin C-432A (**2**) (600 MHz, $CD_3OD$)………………………..…13

**Figure S8.** HSQC spectrum of ketomemicin C-432A (**2**) (600 MHz, $CD_3OD$)……………………..…14

**Figure S9.** HMBC spectrum of ketomemicin C-432A (**2**) (600 MHz, $CD_3OD$)…………………….…15

**Figure S10.** HRMS and MS/MS spectra of ketomemicin C-432A (**2**)………………………………..16

**Table S3.** NMR table for ketomemicin C-432B (**3**)..................................................……………..17

**Figure S11.** [1]H NMR spectrum of ketomemicin C-432B (**3**) (600 MHz, $CD_3OD$)…….………………..18

**Figure S12.** COSY spectrum of ketomemicin C-432B (**3**) (600 MHz, $CD_3OD$)..………………………19

**Figure S13.** HSQC spectrum of ketomemicin C-432B (**3**) (600 MHz, $CD_3OD$).………………………..20

**Figure S14.** HMBC spectrum of ketomemicin C-432B (**3**) (600 MHz, $CD_3OD$).………………………..21

**Figure S15.** HRMS and MS/MS spectra of ketomemicin C-432B (**3**)………..………………………22

**Table S4.** Calculated [13]C NMR chemical shifts for diastereomers of **1**……………...………...……..23

**Table S5.** Calculated [1]H NMR chemical shifts for diastereomers of **1**………………………………..24

**Table S6.** Mean averaged error (MAE) and root-mean-squared deviation (RMSD) between calculated and experimental NMR chemical shifts of **1**………………………………………………………...25

**Table S7.** Summary of DP4+ results between calculated and experimental NMR chemical shifts of **1**.………………………………………………………………………………………………………25

**Figure S16**. Ketomemicin-specific clusters observed via GNPS molecular networking..………...…..26

**Table S4.** Distribution of ketomemicin analogs (**1**–**8**) and ketomemicin biosynthetic gene clusters (*ktm*) in 118 *Salinispora* strains……………………………………..……………………………..27

**Figure S17.** Ketomemicin biosynthetic gene clusters (*ktm*) in *Salinispora* genomes...………………..29

**Figure S18.** Synteny plot of ketomemicin-like biosynthetic gene clusters (*ktm*) in the NCBI database (excluding *Salinispora*)……………………..…………………………………………………30

**Quantum chemical computations.** Conformational searches were conducted using xTB-CREST[1] to explore potential energy surfaces (PES) and identify low-energy conformers for the four possible diastereomers of **1**. To capture energetically favorable conformations, metadynamic simulations were performed at the ALPB(methanol)-GFN2-xTB level with a default step of 5 fs and automatically determined simulation length. Subsequently, each frame in the trajectory file was optimized using the same computational method. Finally, an ensemble of conformers was filtered to group similar conformers using a geometry root-mean-square deviation (RMSD) threshold of 0.5 angstroms and an energy threshold of 0.25 kcal/mol. Conformers within an energy window of 7 kcal/mol were further subjected to DFT optimization. Geometry optimizations and vibrational analyses were conducted using the Gaussian16 Revision C.01 suite. Initially, the IEFPCM(methanol)-RB3LYP-D3(0)/6-31G(d) method was employed to quickly reoptimize the structures from semi-empirical level to DFT level and eliminate duplicated conformers. Structures falling outside the energy window of 7 kcal/mol were discarded. Subsequently, a higher level of theory IEFPCM(methanol)-RB3LYP-D3(0)/6-31+G(d,p) was employed to obtain free energies for subsequent NMR calculations. Only structures within a 3 kcal/mol energy window were selected for the NMR calculation, as a 3 kcal/mol difference corresponds to a Boltzmann population ratio of 0.006 at room temperature (the collection of optimized geometries is available at the ioChem-BD repository via https://iochem-bd.bsc.es/browse/edit-collection/100/316428/8ea6b7f4c36295411c31f717. Conformers were subjected to NMR calculations using the IEFPCM(methanol)-mpw1pw91/ 6-311+G(2d,p) level of theory. The isotropic values obtained from these calculations were converted to predicted shifts using the scaling factors provided by CHESHIRE (http://cheshirenmr.info/ScalingFactors.htm).[2] The slope and intercept factors for $^{1}$H are -1.0754 and 31.8463 ppm, while for $^{13}$C these are -1.0399 and 186.5993 ppm. The computed shifts were subsequently averaged based on the Boltzmann weight of each conformer. Chemical shifts for equivalent hydrogen and carbon atoms were averaged. Experimental and computed chemical shift values for each diastereomer were compared using the mean averaged error (MAE), linear coefficient of determination $R^2$, root-mean-squared deviation (RMSD), and DP4+ methods.[3] The DP4+ analysis was conducted following the instructions and Excel table available at http://www.sarotti-NMR.weebly.com or http://www.iquir-conicet.gov.ar/Sarotti-NMR.


1. Pracht, P., Bohle, F., & Grimme, S. (2020). *Physical Chemistry Chemical Physics*, *22*(14), 7169–7192. doi: 10.1039/C9CP06869D.
2. Lodewyk, M. W., Siebert, M. R., & Tantillo, D. J. (2012). *Chemical Reviews*, 112, 1839-1862. doi: 10.1021/cr200106v.
3. Zanardi, M. M., & Sarotti, A. M. (2021). *The Journal of Organic Chemistry*, *86*(12), 8544–8548. doi: 10.1021/acs.joc.1c00987.


***Salinispora* genome and *ktm* BGC phylogeny.** The *Salinispora* genome phylogeny used here was previously described by Román-Ponce et al. 2020. For the *ktm* BGC phylogeny we first we identified 79 *Salinispora* strains with the six-gene *ktm* using Cblaster. The translated amino acid sequences of *ktm A-F* were then extracted, concatenated, and aligned using MUSCLE. The alignment was analyzed in raxmlGUI 2.0,[4] using BLOSUM62 matrix, ML + rapid boodstrap (1000 repetitions), and GAMMA substitution rates. The resulting trees were visualized and edited using iTOL v5.[5]


4. Letunic I. and Bork P. (2021) *Nucleic Acids Research*, 49, W293-W296. doi: 10.1093/nar/gkab301
5. Edler, D., Klein, J., Antonelli, A., and Silvestro, D. (2020) *Methods in Ecology and Evolution*, doi: 10.1111/2041-210X.13512.


**Protease inhibition studies.** SARS-CoV-2 $M^{pro}$ and cruzain were recombinantly expressed as described previously.[6,7] Recombinant *Trypanosoma brucei* cathepsin L (TbrCatL) was generously provided by Conor Caffrey (University of California San Diego).[8] Other recombinant proteases SARS-CoV-1 3CL/M$^{pro}$ (E-718), MERS-CoV 3CL/M$^{pro}$ (E-719), human cathepsin B (953-CY), human cathepsin L (#952-CY-010), human cathepsin D (1014-AS), human aminopeptidase B (8089-ZN-010), human 20S proteasome protein (h20S; E-360-050), and recombinant human PA28 activator alpha subunit (PA28; E-381-100) were purchased from R&D Systems, while SARS-CoV-2 PL$^{pro}$ was purchased from Acro Biosystems (PAE-C518). The final enzyme concentration in each assay was 0.5 nM for cruzain, TbrCatL, c20S, and PA28; 25 pM for human cathepsin L; 50 nM for SARS-CoV, SARS-CoV-2 M$^{pro}$, MERS-CoV M$^{pro}$, and SARS-CoV-2 PL$^{pro}$; 8 ng/µL for human cathepsin D; 675 pM for human aminopeptidase B; and 0.1 ng/µL for Cathepsin B. Fluorogenic peptide substrates were used at the following concentrations: 10 µM of Ac-Abu-Tle-Leu-Gln-AMC (Vivitide, SFP-3250-v) for SARS-CoV M$^{pro}$, SARS-CoV-2 M$^{pro}$, and MERS-CoV M$^{pro}$; 2.5 µM of z-Phe-Arg-AMC (R&D Systems, ES009) for cruzain and TbrCatL, 25 µM of z-Phe-Arg-AMC (R&D Systems, ES009) for human cathepsin B and L; 5 µM of Mca-Gly-Lys-Pro-Ile-Leu-Phe-Phe-Arg-Leu-Lys(DNP)-dArg-NH2 (CPC Scientific, SUBS-017A) for human cathepsin D; 100 µM of Arg-AMC (Bachem I1050) for human aminopeptidase B; 50 µM of Suc-Arg-Leu-Arg-Gly-Gly-AMC for SARS-CoV-2 PLpro; 50 µM of Suc-Leu-Leu-Val-Tyr-AMC (Bachem 4011369) for human 20S proteasome β5 subunit; z-Leu-Leu-Glu-AMC (R&D Systems S230) for human 20S proteasome β1 subunit, and Boc-Leu-Arg-Arg-AMC (AdipoGen Life Sciences AG-CP3-0014) for human 20S proteasome β2 subunit. Assay buffers for protease assays were as follows, Assay Buffer 1 (SARS-CoV M$^{pro}$, SARS-CoV-2 M$^{pro}$, and MERS-CoV M$^{pro}$): 50 mM HEPES pH7.5, 150 mM NaCl, 1 mM EDTA, 0.01% Tween 20; Assay Buffer 2 (Cruzain and TbrCatL): 100 mM Na-Acetate pH 5.5, 1 mM DTT, 0.01 % Triton X-100; Assay Buffer 3 (Cathepsin B and Cathepsin L): 50 mM Na-Acetate pH 5.5, 5 mM DTT, 1 mM EDTA, 100 mM NaCl, 0.01% Triton X-100 (0.01 % BSA was also added to cathepsin L assay); Assay Buffer 4 (Cathepsin D): 100 mM Citrate-Phosphate, pH 3.6; Assay Buffer 5 (human aminopeptidase B): 50 mM Tris pH 7.5, 100 mM KCl, 1 mM DTT; Assay Buffer 6 (human 20S proteasome): 50 mM HEPES pH7.5 and 1 mM DTT, and Assay Buffer 7 (SARS-CoV-2 PL$^{pro}$): 50 mM HEPES pH 6.5, 150 mM NaCl, 0.01% Tween 20, 0.1 mM DTT. Assays were performed with 10 µM of the ketomemicins. Control inhibitors include 10 µM E-64 (Sigma E3132), 10 µM Pepstatin (Sigma P5318), 100 nM Nirmatrelvir (Selleckchem S9866), 10 µM Bestatin (APEXBio A8621), 10 µM Salinosporamide A (Marizomib, Sigma SML1916), and 10 µM GRL-0617 (Sigma SML2961). The compound was diluted into the appropriate assay buffer and pre-incubated with protease for 15 minutes at 25°C. Human 20S proteasome was incubated for 1h with 10 molar equivalents of PA28 activator before the pre-incubation with the ketomemicins for additional 30 minutes. The addition of substrate in the assay buffer initiated the reaction. All assays were performed at 25°C in triplicate wells with DMSO as the vehicle control. The final volume of each reaction was 30 µL in a 384-well black plate. Fluorescence was measured at 360/460 nm (ex/em) for peptide-AMC substrates or 320/400 nm (ex/em) for internally quenched MCA-peptide-DNP substrates in a Biotek Synergy HTX fluorescence plate reader. Enzymatic activity was calculated by comparison to the initial rates of reaction of a DMSO control.

6. Mellott, D. M., Tseng, C.-T., Drelich, A., Fajtová, P., *et al.* (2021). *ACS Chemical Biology*, *16*(4), 642–650. https://doi.org/10.1021/acschembio.0c00875
7. Barbosa da Silva, E., Dall, E., Briza, P., Brandstetter, H., *et al.* (2019). *Acta Crystallographica Section F Structural Biology Communications*, *75*(6), 419–427. https://doi.org/10.1107/S2053230X19006320
8. Caffrey CR, Hansell E, Lucas KD, Brinen LS, Alvarez Hernandez A, Cheng J, Gwaltney SL 2nd, Roush WR, Stierhof YD, Bogyo M, Steverding D, McKerrow JH. Mol Biochem Parasitol. 2001 Nov;118(1):61-73. doi: 10.1016/s0166-6851(01)00368-1. PMID: 11704274.

**Table S1.** NMR table for ketomemicin C-418 (**1**) (600 MHz, $CD_3OD$)

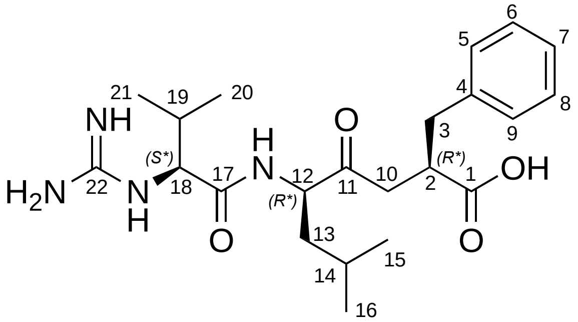


| position | $\delta_H$, mult. ($J$ in Hz) | $\delta_C$[a], type | COSY | HMBC |
|---|---|---|---|---|
| 1 | ---- | 182.0, C | ---- | ---- |
| 2 | 3.01 | 46.4, CH | 3a, 3b, 10 | |
| 3 | a 3.06, dd (13.5, 5.0)<br>b 2.65, dd (12.5, 4.0) | 39.6, $CH_2$ | 2, 3b<br>2, 3a | 1, 2, 4, 5, 9, 10<br>1, 2, 4, 5, 9, 10 |
| 4 | ---- | 141.5, C | ---- | ---- |
| 5, 9 | 7.23, d (6.6) | 130.0, CH | 6, 8 | 3, 4, 5, 7, 9 |
| 6, 8 | 7.25, t (7.5) | 129.1, CH | 5, 7, 9 | 4, 6, 8 |
| 7 | 7.16 (t, 6.7) | 126.9, CH | 6, 8 | 4, 5, 9 |
| 10 | a 2.87, dd (16.8, 9.4)<br>b 2.32, d (17.2) | 41.7, $CH_2$ | 2 | 1, 2, 3, 11 |
| 11 | ---- | 209.4, C | ---- | ---- |
| 12 | 4.43, dd (11.1, 2.8) | 58.5, CH | 13a, 13b | 11, 13, 14, 17 |
| 13 | a 1.67, m (10.5)<br>b 1.47, dd (10.9, 3.5) | 39.7, $CH_2$ | 13b<br>13a | 12, 14, 15, 16 |
| 14 | 1.63, bs | 26.0, CH | 15, 16 | |
| 15 | 0.94, d (6.3) | 23.6, $CH_3$ | | 13, 14, 16 |
| 16 | 0.92, d (6.3) | 21.4, $CH_3$ | | 13, 14, 15 |
| 17 | ---- | 171.7, C | ---- | ---- |
| 18 | 3.85, d (6.3) | 62.0, CH | 19, 20 | 17, 19, 20, 21, 22 |
| 19 | 2.19, sex (6.6) | 32.1, CH | 18, 20 | |
| 20 | 0.98, d (6.9) | 19.4, $CH_3$ | | 18, 19, 21 |
| 21 | 0.95, d (6.9) | 17.8, $CH_3$ | | 18, 19, 20 |
| 22 | ---- | 158.9, C | ---- | ---- |

[a] Carbon assignments were made on the basis of HSQC and HMBC experiments

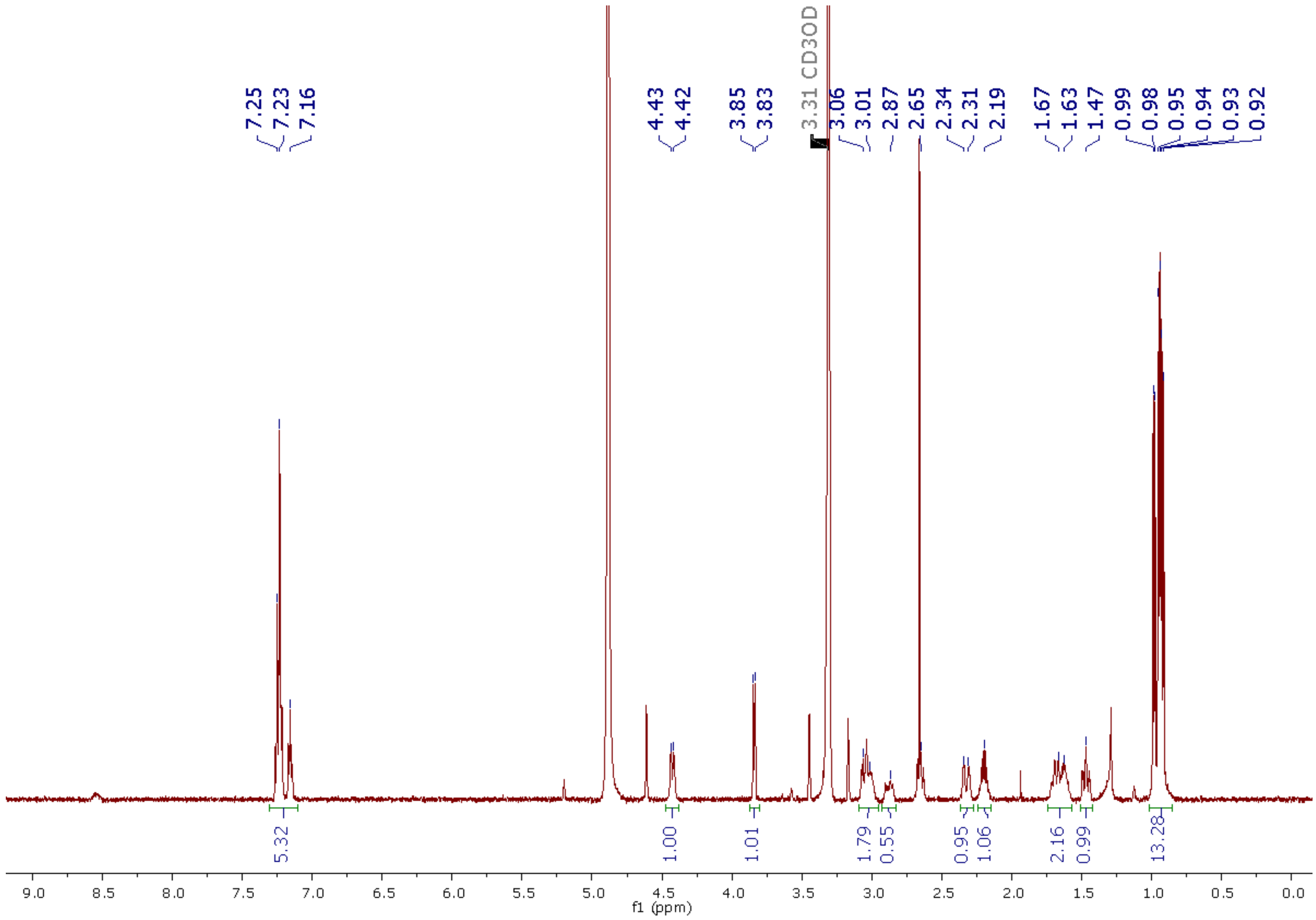


**Figure S1.** $^1$H NMR spectrum of ketomemicin C-418 (**1**) (600 MHz, $CD_3OD$)

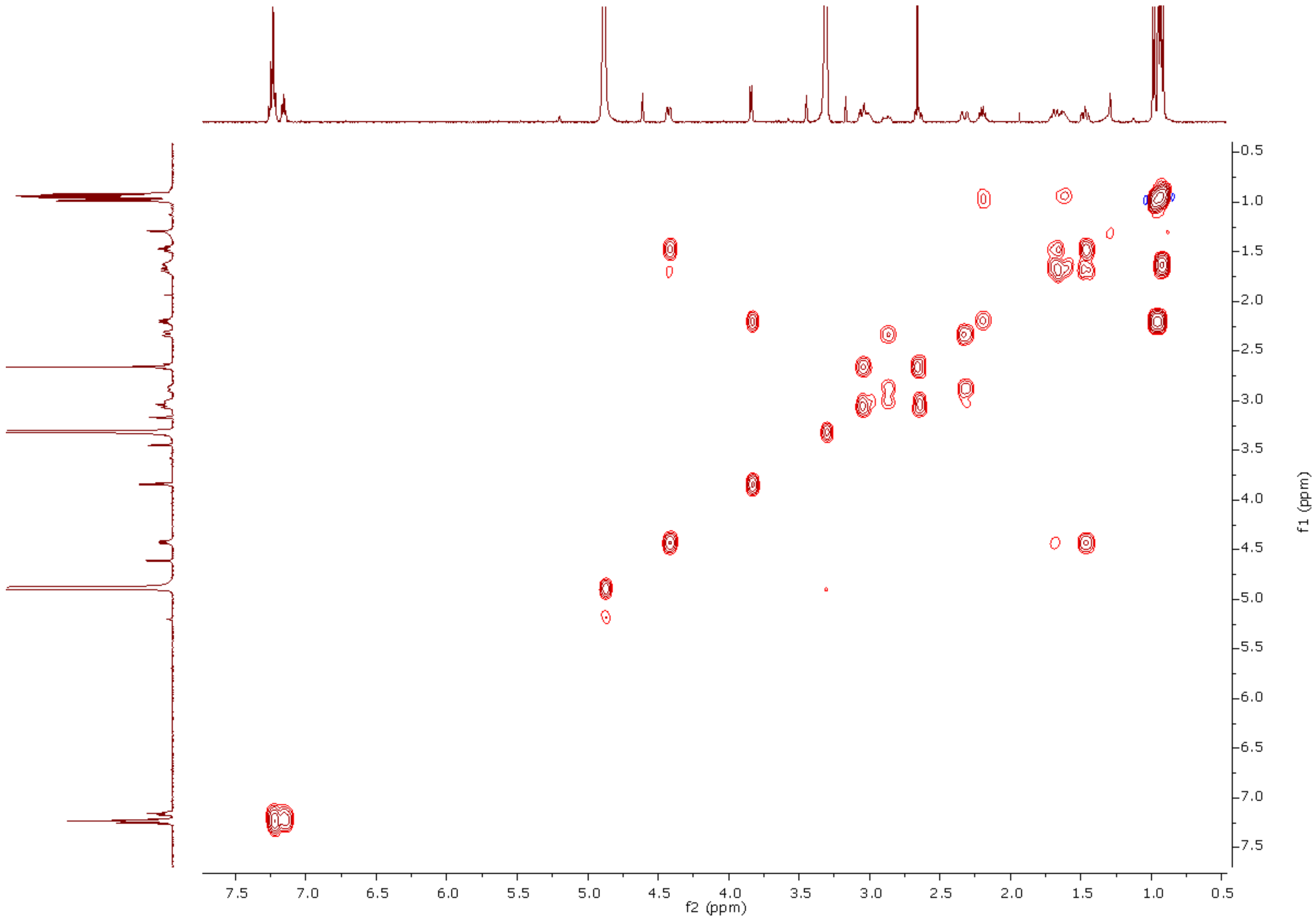


**Figure S2.** COSY spectrum of ketomemicin C-418 (**1**) (600 MHz, $CD_3OD$)

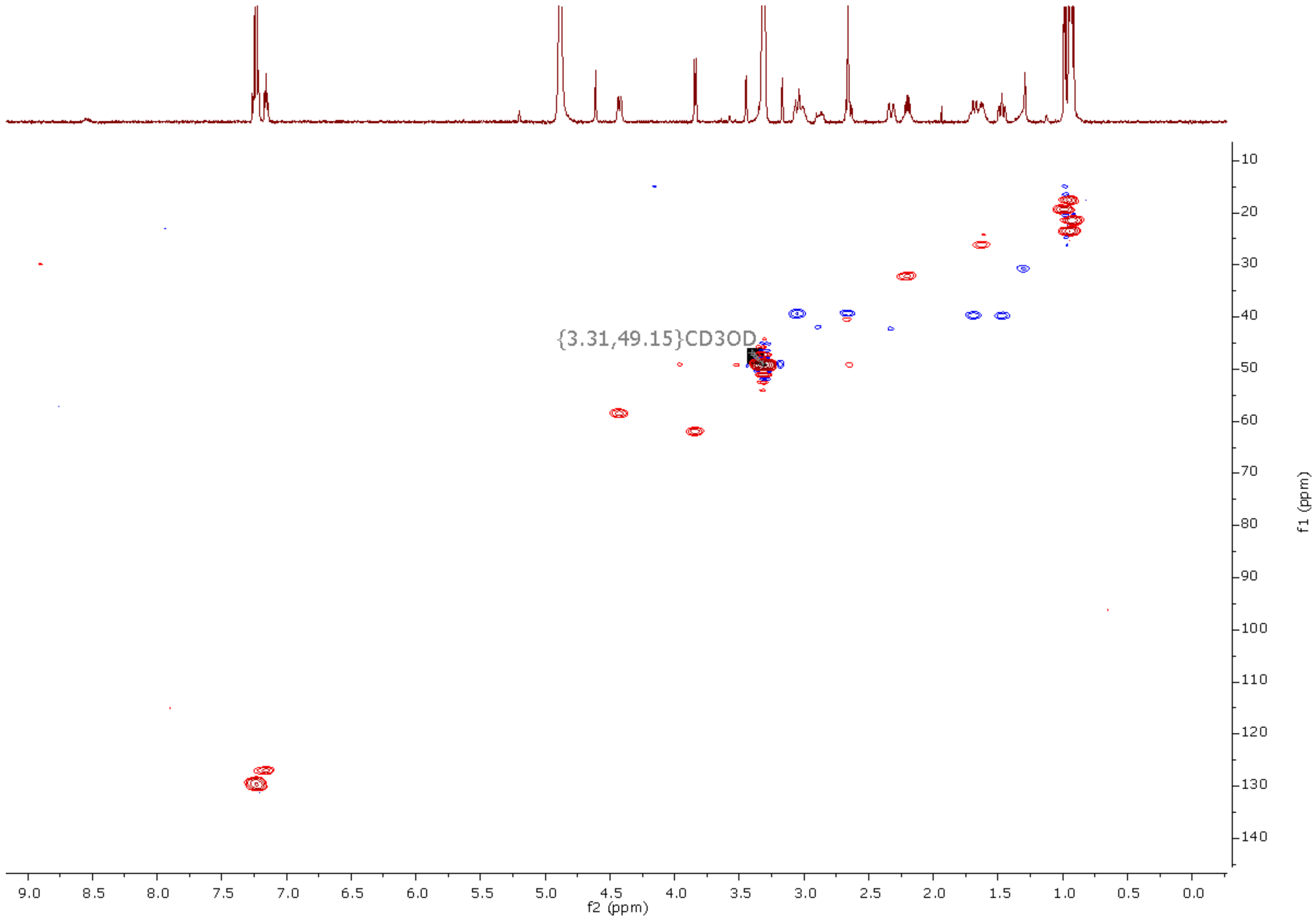


**Figure S3.** HSQC spectrum of ketomemicin C-418 (**1**) (600 MHz, $CD_3OD$)

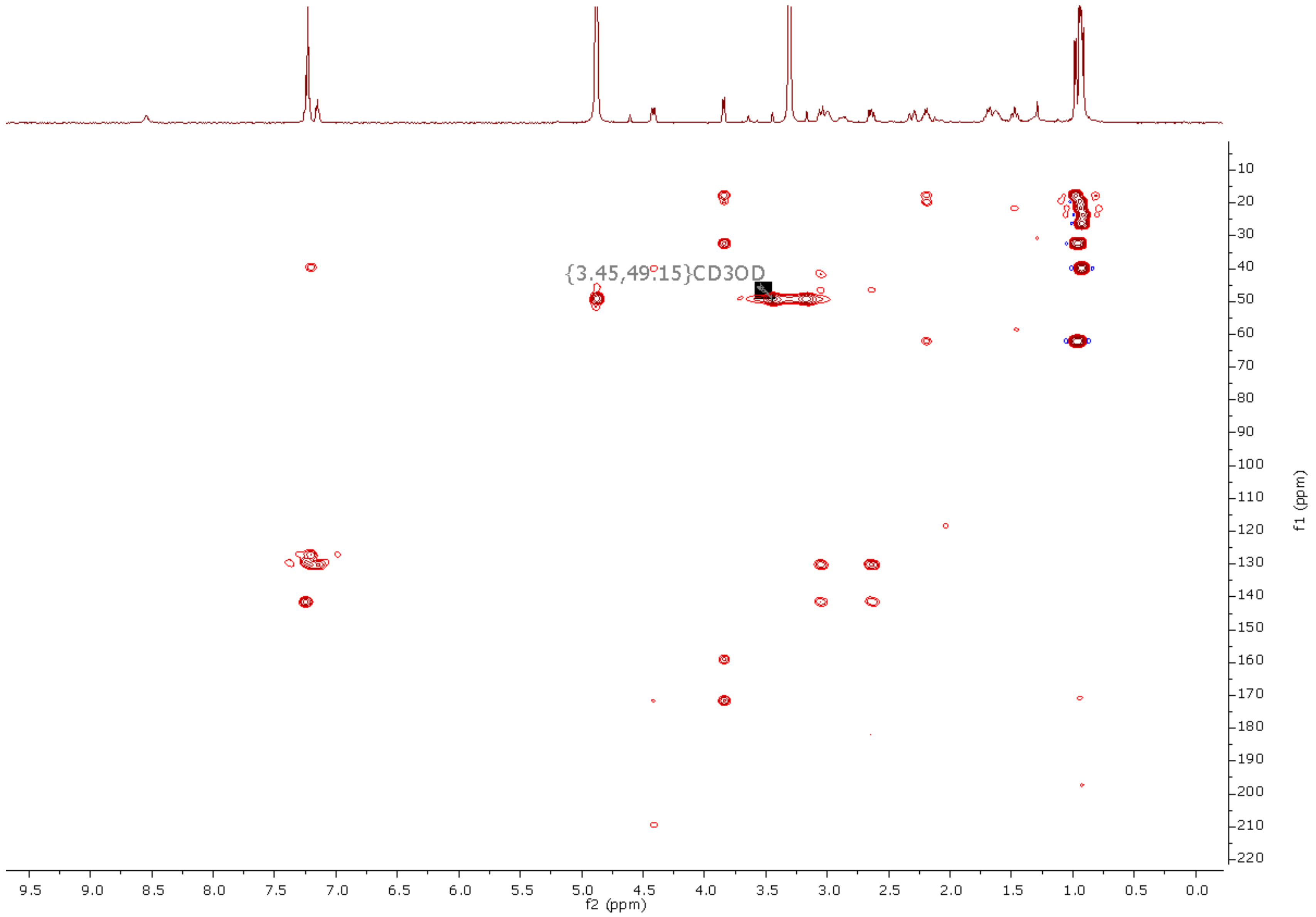


**Figure S4.** HMBC spectrum of ketomemicin C-418 (**1**) (600 MHz, $CD_3OD$)

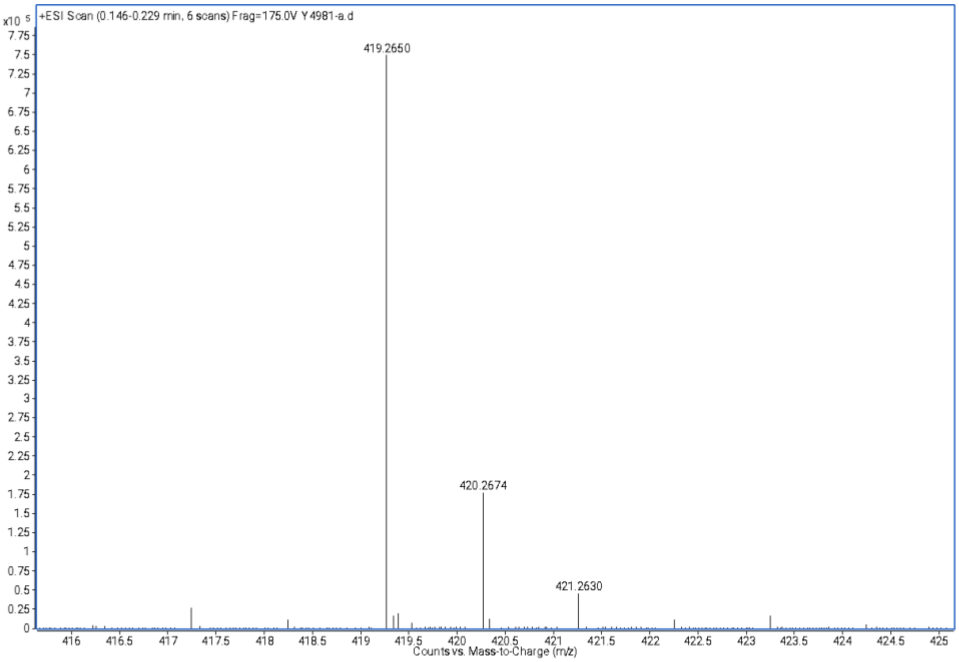


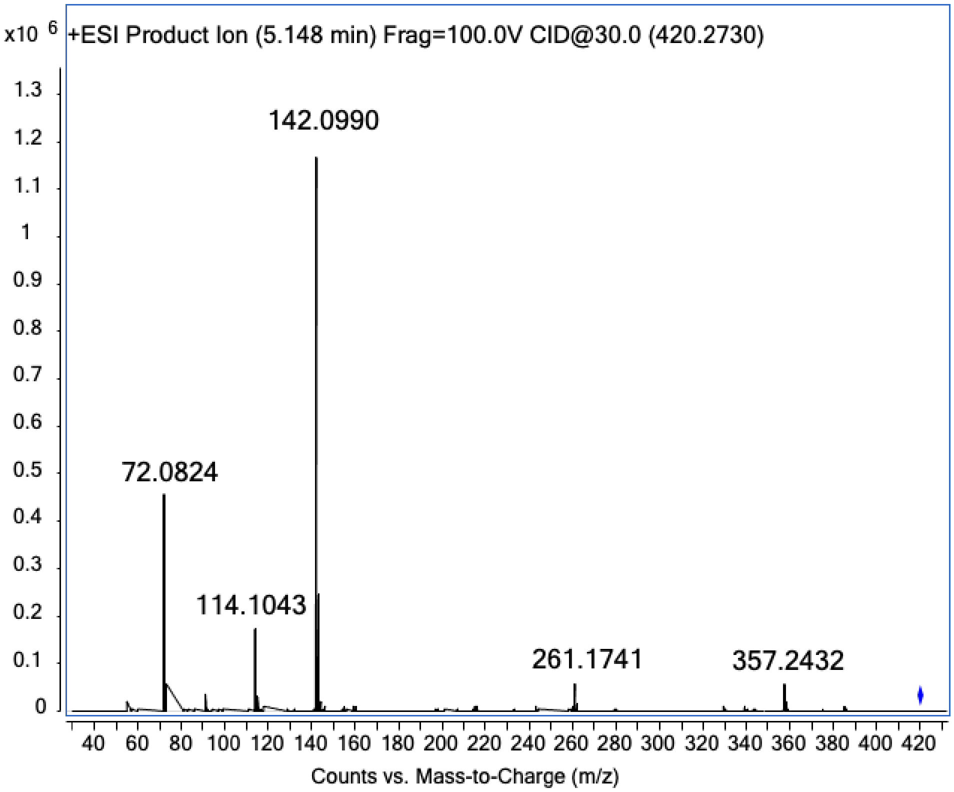


**Figure S5.** HRMS and MS/MS spectra of ketomemicin C-418 (**1**)

**Table S2.** NMR table for ketomemicin C-432A (**2**) (600 MHz, $CD_3OD$)

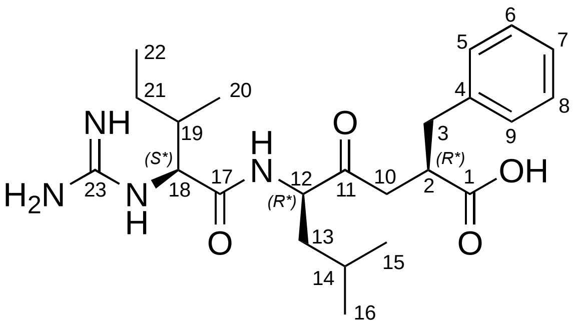


| position | $\delta_H$, mult. (*J* in Hz) | $\delta_C$[a], type | COSY | HMBC |
|---|---|---|---|---|
| 1 | ---- | 181.8, C | ---- | ---- |
| 2 | 3.03 [b] | 46.4, CH | | |
| 3 | a 3.04, m<br>b 2.66, m | 39.4, $CH_2$ | 3b<br>3a | 2, 4, 5, 9, 10<br>2, 4, 5, 9, 10 |
| 4 | ---- | 141.2, C | ---- | ---- |
| 5, 9 | 7.23, d (6.9) | 130.4, CH | 6, 8 | 3, 5, 7, 9 |
| 6, 8 | 7.25, t (7.1) | 129.2, CH | 5, 7, 9 | 4, 6, 8 |
| 7 | 7.16, t (7.0) | 127.1, CH | 6, 8 | 5, 9 |
| 10 | a 2.86, m<br>b 2.35, m | 42.0, $CH_2$ | 10b<br>10a | 3 |
| 11 | ---- | 209.3, C | ---- | ---- |
| 12 | 4.43, d (10.2) | 58.4, CH | 13b | 13, 14 |
| 13 | a 1.65 [b]<br>b 1.48, m | 39.6, $CH_2$ | 12, 13b<br>12, 13a, 14 | 11, 15, 16<br>12, 15, 16 |
| 14 | 1.64 [b] | 26.0, CH | 15, 16 | |
| 15 | 0.93 [b] | 23.5, $CH_3$ | 14 | 13, 14, 16 |
| 16 | 0.91 [b] | 21.4, $CH_3$ | 14 | 13, 14, 15 |
| 17 | ---- | 171.8, C | ---- | ---- |
| 18 | 3.91, d (6.7) | 61.3, CH | 19, 20 | 17, 19, 20, 21, 23 |
| 19 | 1.94, m | 38.4, CH | 18, 20, 21a, 21b | 18 |
| 20 | 0.96 [b] | 15.6, $CH_3$ | 19 | 18, 19, 21 |
| 21 | a 1.52 [b]<br>b 1.17, m | 25.3, CH | 21b<br>21a | <br>19, 20 |
| 22 | 0.93 [b] | 11.4, $CH_3$ | ---- | ---- |
| 23 | ---- | 158.8, C |  |  |

[a] Carbon assignments were made on the basis of HSQC and HMBC experiments

[b] Signal partially obscured

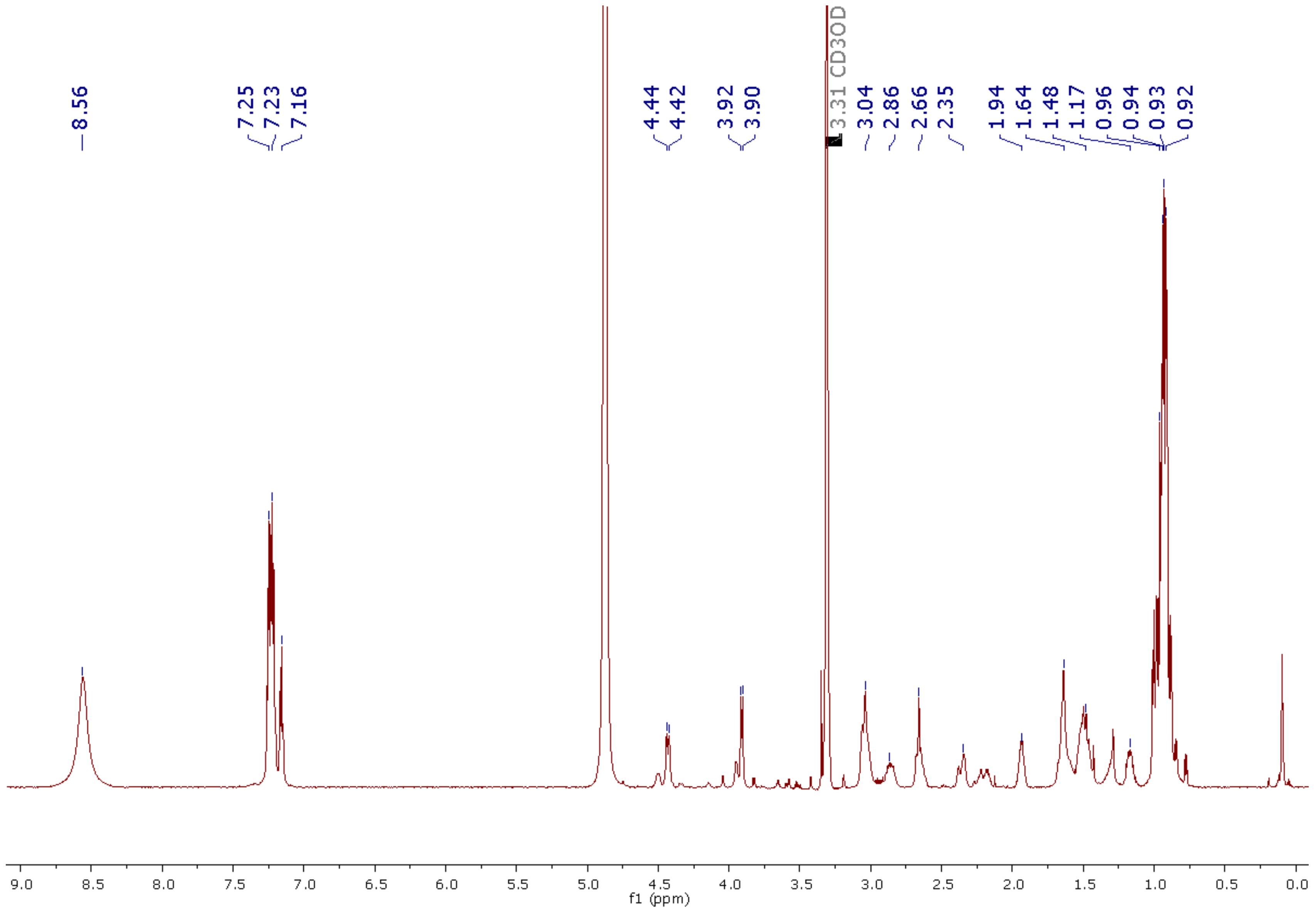


**Figure S6.** 1H NMR spectrum of ketomemicin C-432A (**2**) (600 MHz, $CD_3OD$)

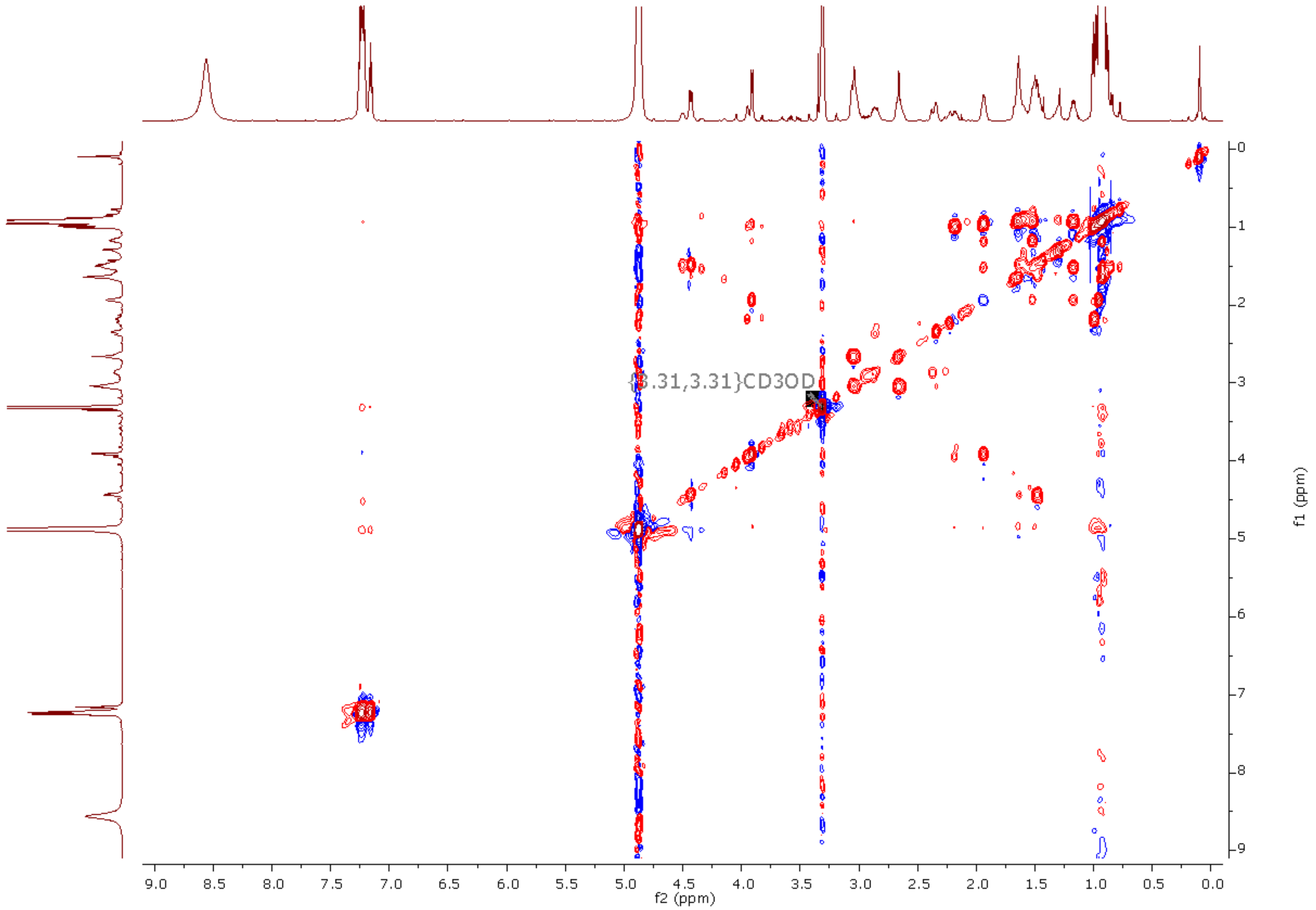


**Figure S7**. COSY spectrum of ketomemicin C-432A (**2**) (600 MHz, $CD_3OD$)

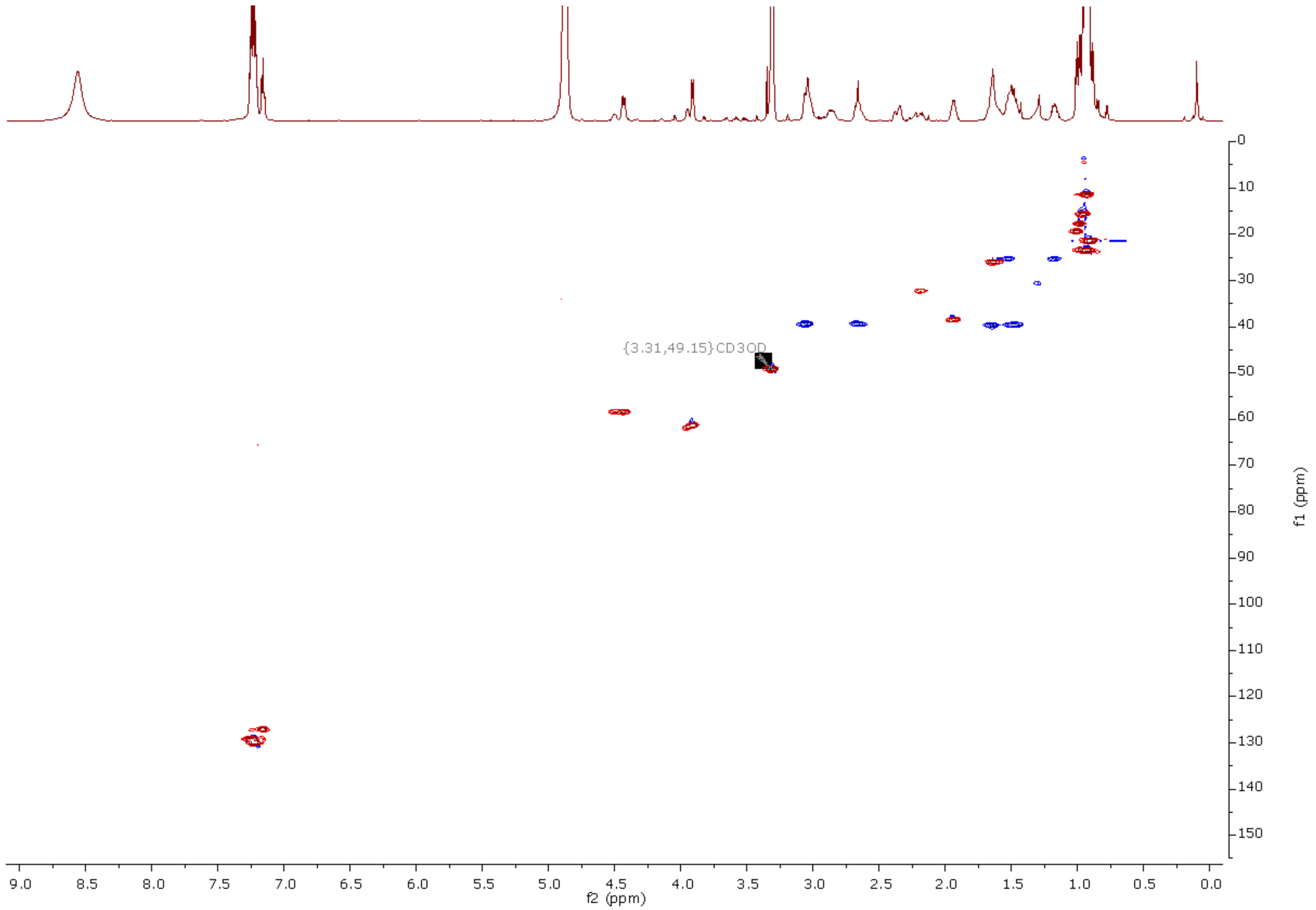


**Figure S8**. HSQC spectrum of ketomemicin C-432A (**2**) (600 MHz, $CD_3OD$)

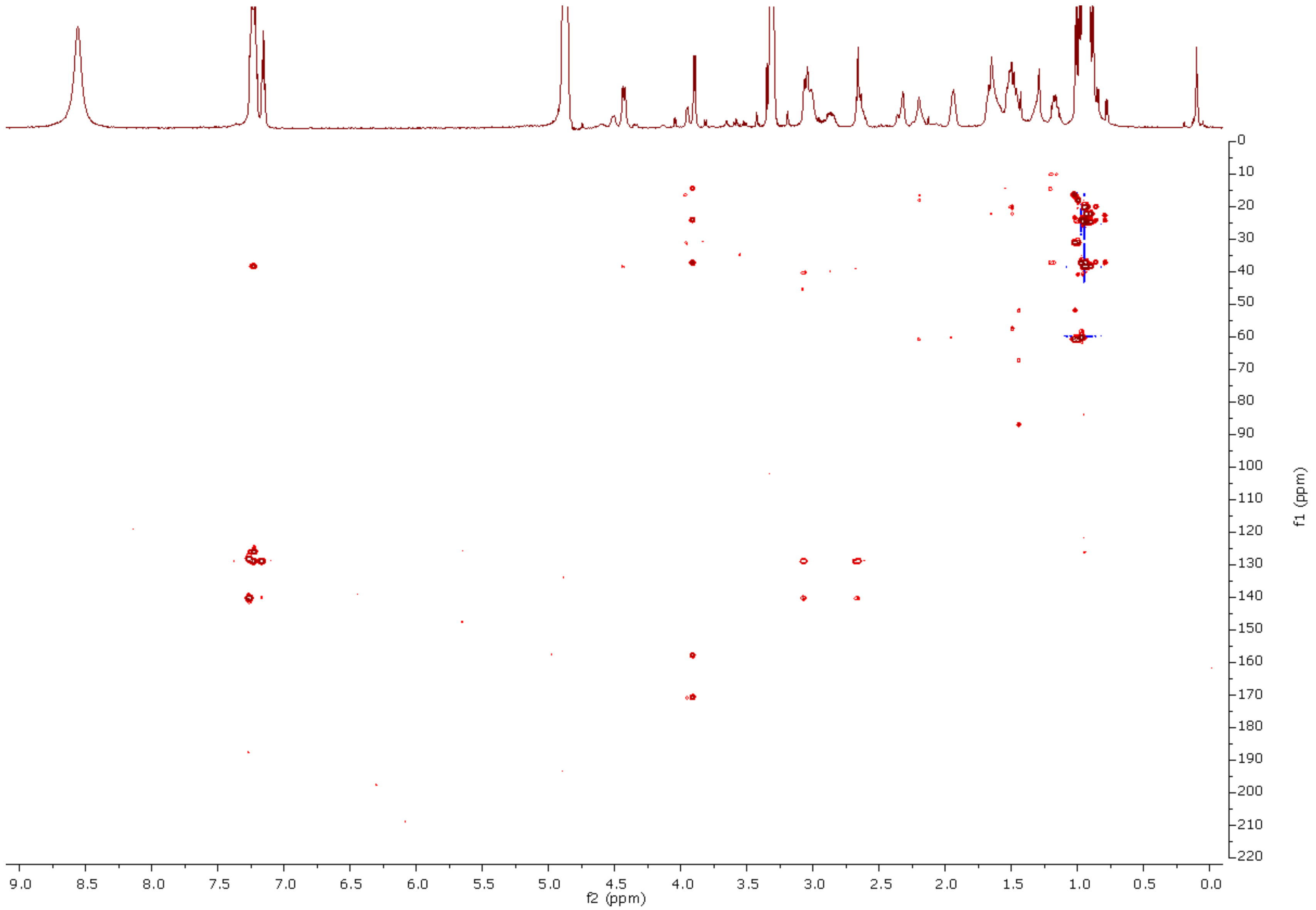


**Figure S9**. HMBC spectrum of ketomemicin C-432A (**2**) (600 MHz, $CD_3OD$)

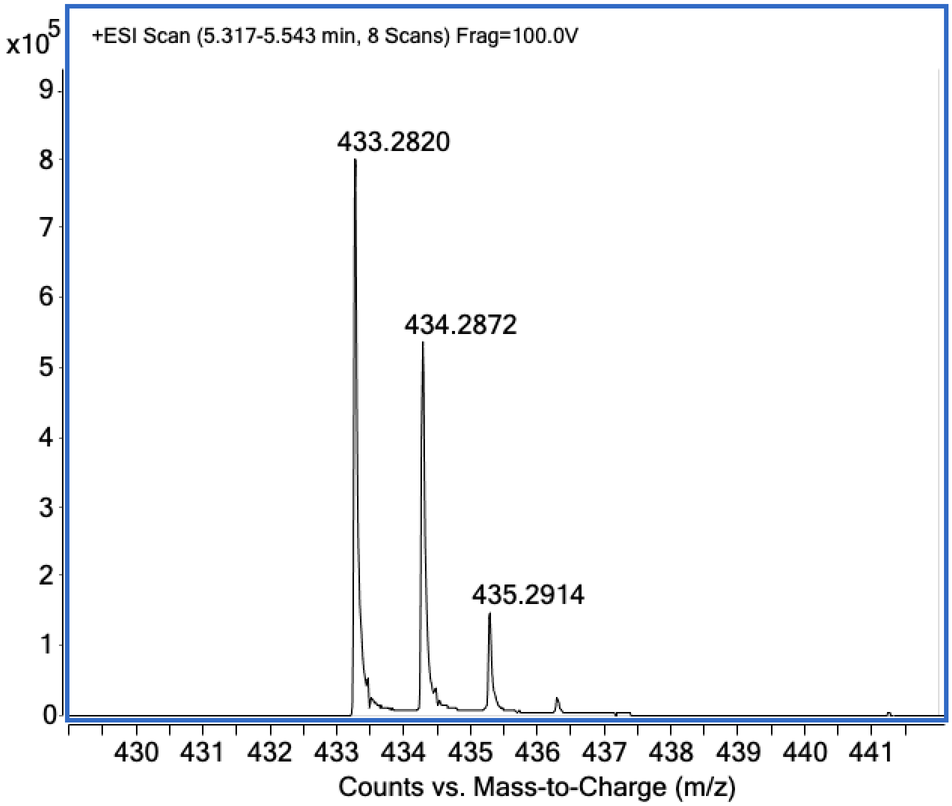


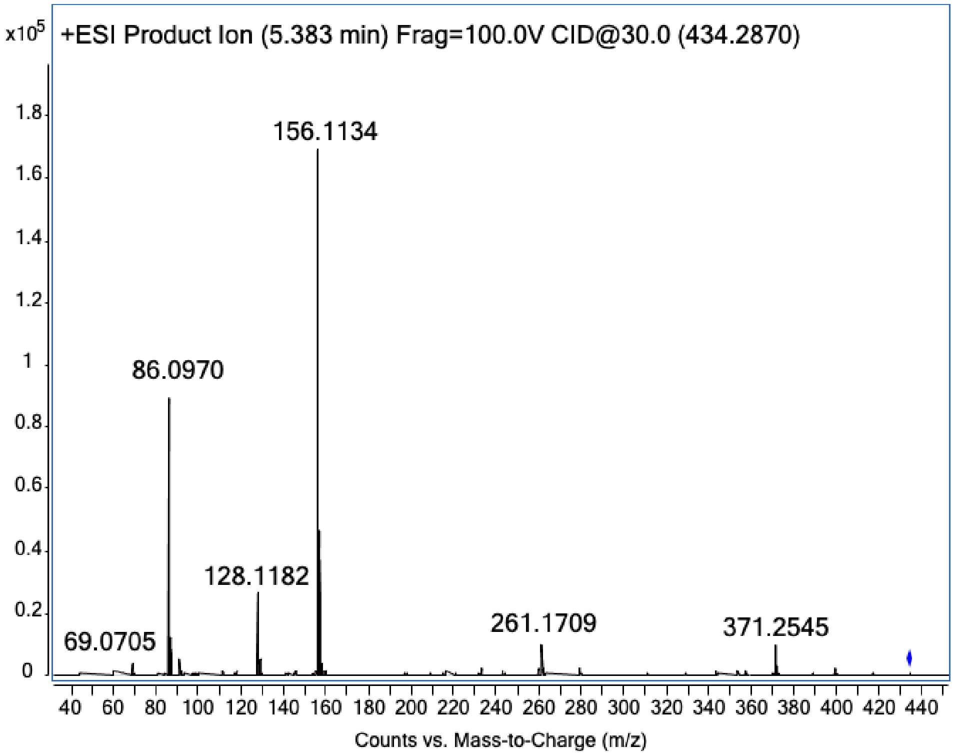


**Figure S10**. HRMS and MS/MS spectra of ketomemicin C-432A (**2**)

**Table S3.** NMR table for ketomemicin C-432B (**3**) (500 MHz, $CD_3OD$)

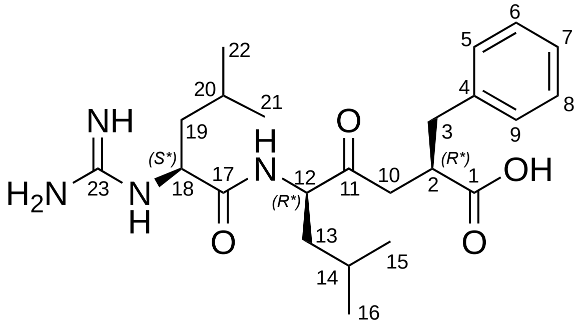

| c | $\delta_H$, mult. ($J$ in Hz) | $\delta_C$[a], type | COSY | HMBC |
|---|---|---|---|---|
| 1 | ---- | 182.0, C | ---- | ---- |
| 2 | 3.00, bm | 46.2, CH | 3a, 3b, 10 | |
| 3 | a 3.05, dd (13.3, 5.2)<br>b 2.65, dd (13.0, 8.9) | 39.4, $CH_2$ | 2, 3b, 10<br>3a, 10 | 1, 2, 4, 5, 9, 10<br>1, 2, 4, 5, 9, 10 |
| 4 | ---- | 141.4, C | ---- | ---- |
| 5, 9 | 7.22, d (7.4) | 130.0, CH | 6, 8 | 3, 5, 7, 9 |
| 6, 8 | 7.25, t (7.2) | 129.2, CH | 5, 7, 9 | 4, 6, 8 |
| 7 | 7.15, t (7.2) | 127.0, CH | 6, 8 | 5, 9 |
| 10 | a 2.86 [b]<br>b 2.31, m | 41.5, $CH_2$ | 10b [c]<br>2, 10a [c] | <br>1, 2, 3, 11 |
| 11 | ---- | 209.6, C | ---- | ---- |
| 12 | 4.39, dd (11.1, 3.0) | 58.5, CH | 13a, 13b | 11, 13, 14, 17 |
| 13 | a 1.66 [b]<br>b 1.48, dt | 39.6, $CH_2$ | 12, 13b<br>12, 13a, 14 | 12, 14, 16<br>12, 14, 16 |
| 14 | 1.63 [b] | 26.0, CH | 13, 15, 16 | |
| 15 | 0.93 [b] | 23.5, $CH_3$ | 14 | 13, 14, 16 |
| 16 | 0.91, d (5.9) | 21.4, $CH_3$ | 14 | 13, 14, 15 |
| 17 | ---- | 172.6, C | ---- | ---- |
| 18 | 4.12, dd (9.2, 4.0) | 55.3, CH | 19 | 17, 19, 20, 23 |
| 19 | 1.64 [b] | 42.0, $CH_2$ | 18, 20 | 17, 18 |
| 20 | 1.68 [b] | 25.8, CH | 19, 21, 22 | 19 |
| 21 | 0.94 [b] | 21.7, $CH_3$ | 20 | 19, 20, 22 |
| 22 | 0.98, d (5.8) | 23.2, $CH_3$ | 20 | 19, 20, 21 |
| 23 | ---- | 158.7, C | ---- | ---- |

[a] Carbon assignments were made on the basis of HSQC and HMBC experiments

[b] Signal partially obscured

[c] Denotes weak signal

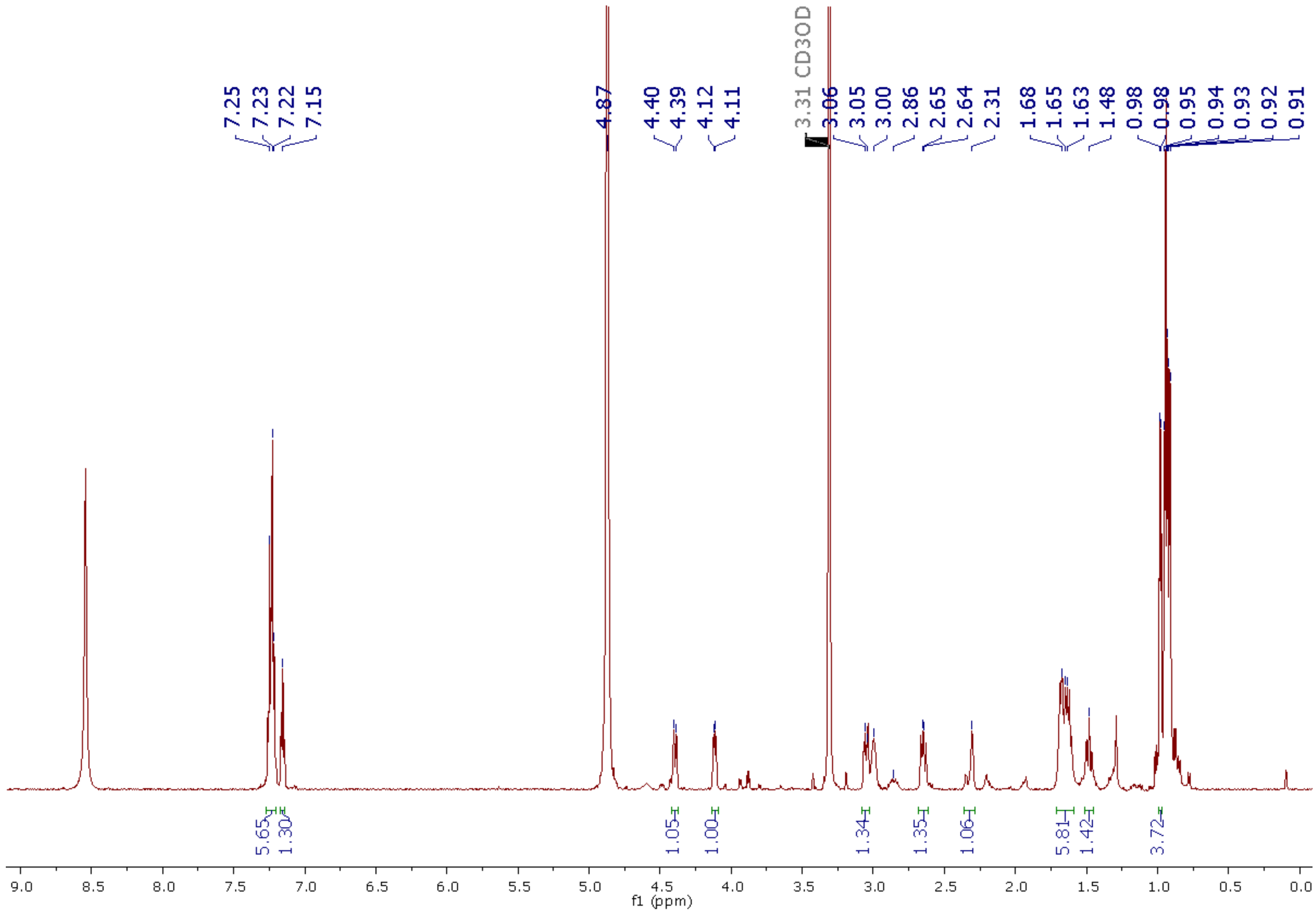


**Figure S11.** 1H NMR spectrum of ketomemicin C-432B (**3**) (600 MHz, $CD_3OD$)

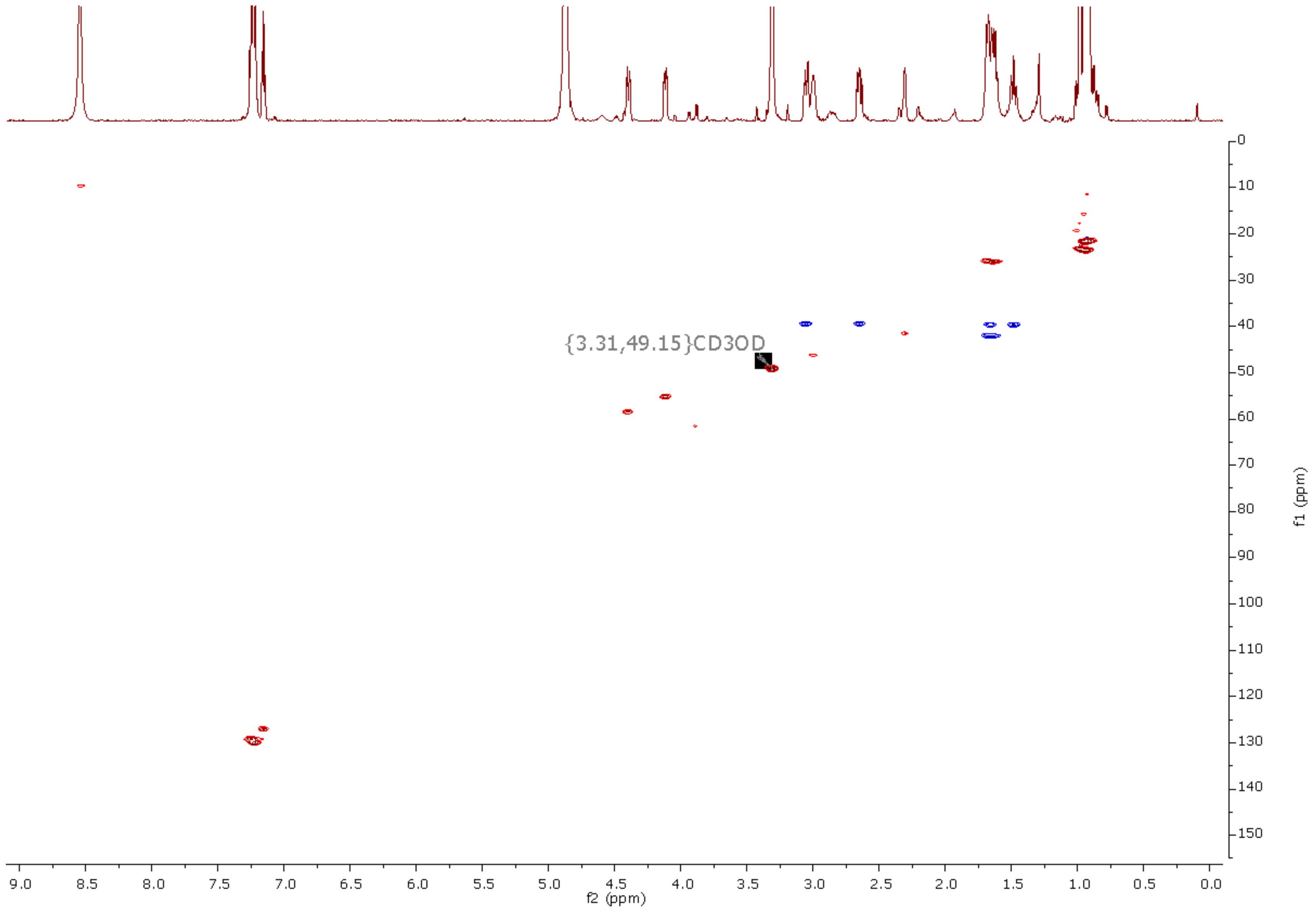


**Figure S12.** HSQC spectrum of ketomemicin C-432B (**3**) (600 MHz, $CD_3OD$)

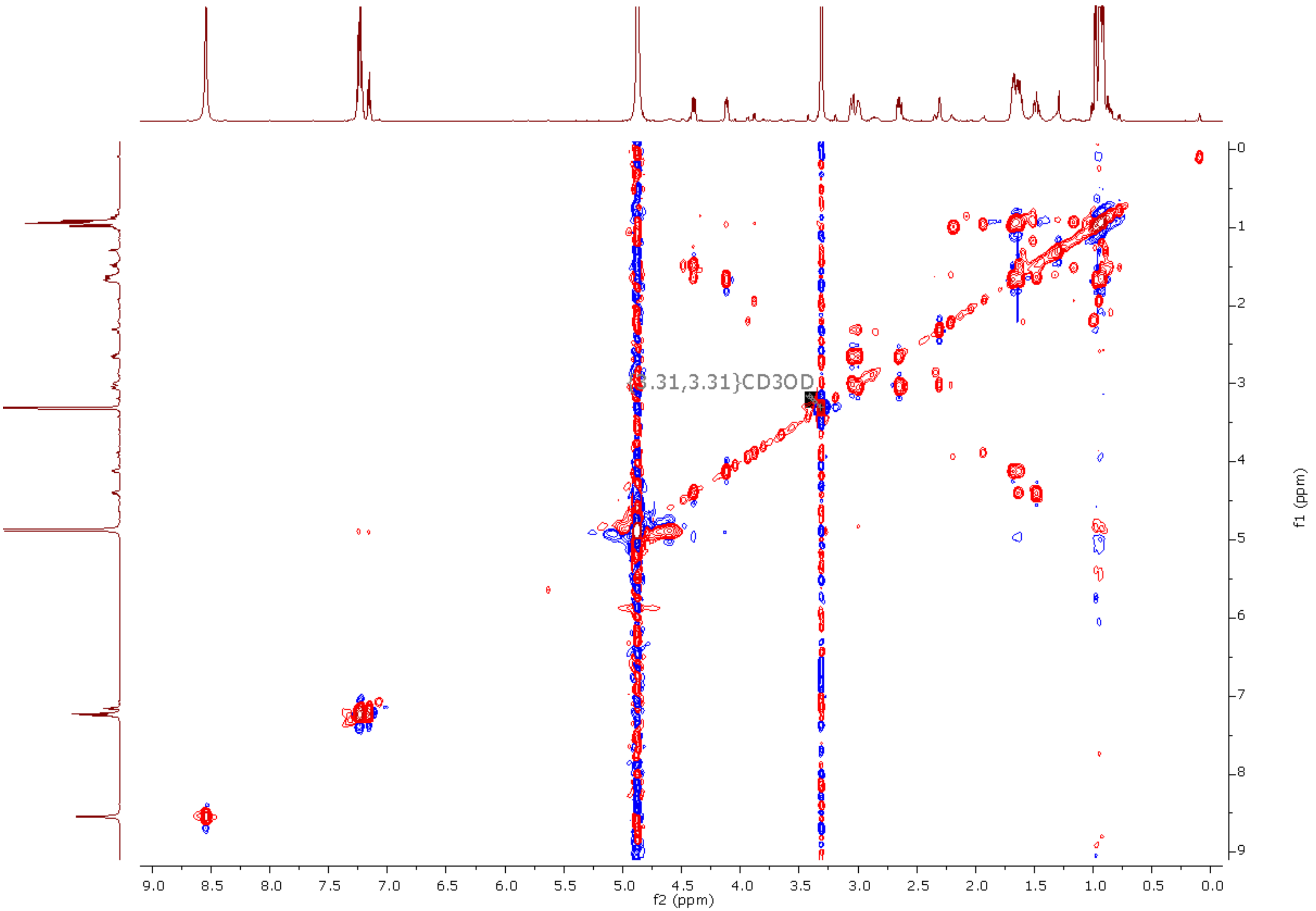


**Figure S13.** COSY spectrum of ketomemicin C-432B (**3**) (600 MHz, $CD_3OD$)

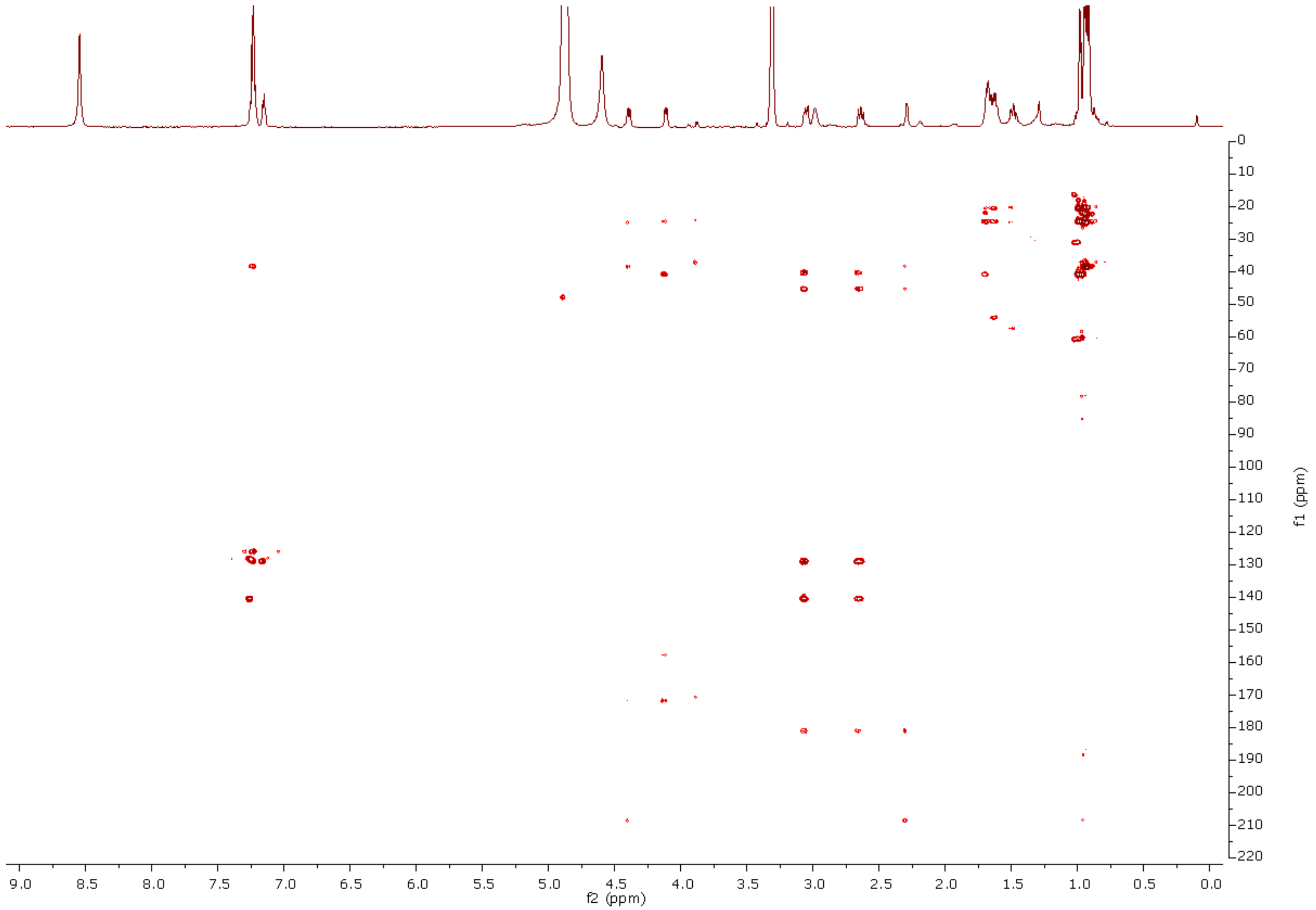


**Figure S14.** HMBC spectrum of ketomemicin C-432B (**3**) (600 MHz, $CD_3OD$)

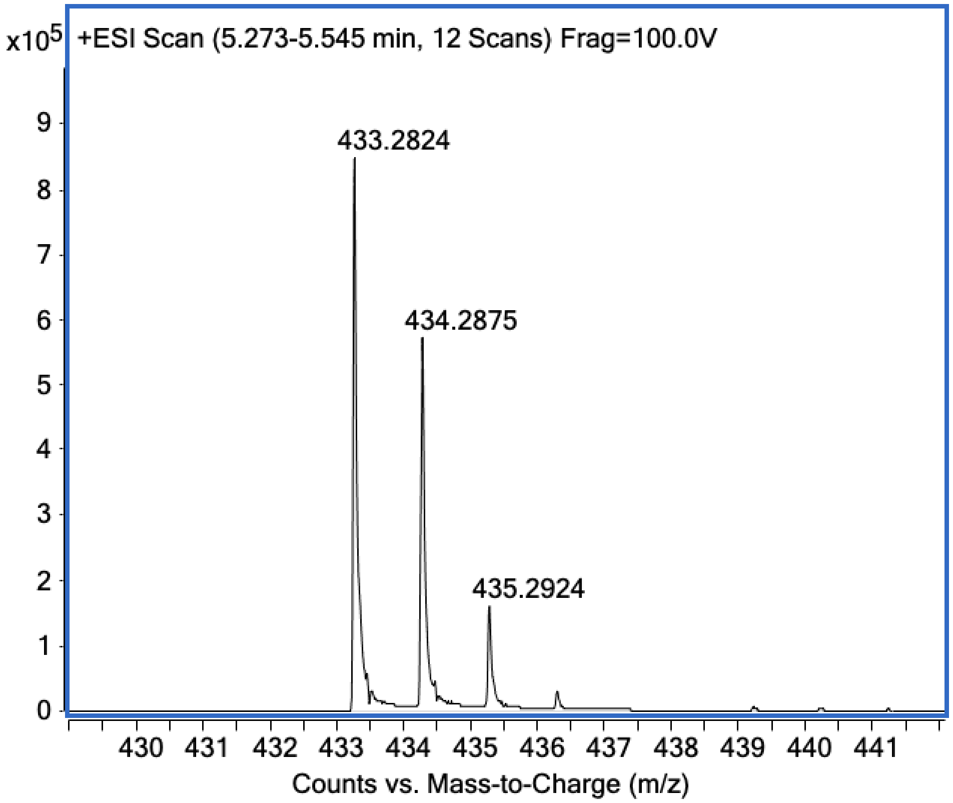


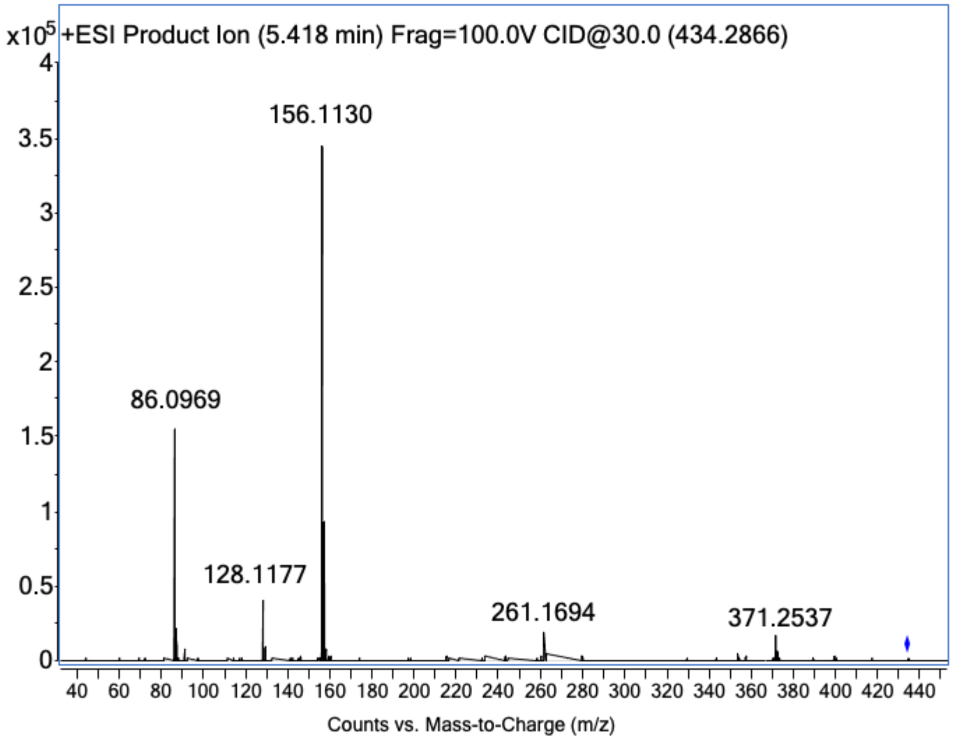


**Figure S15**. HRMS and MS/MS spectra of ketomemicin C-432B (**3**)

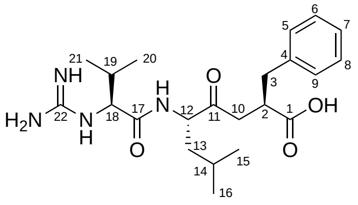

2*R**, 12*S**, 18*S**-**1**

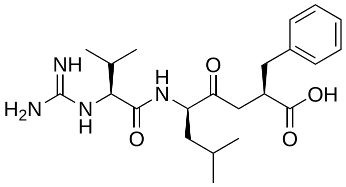

2*R**, 12*R**, 18*S**-**1**

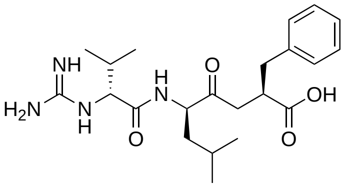

2*R**, 12*R**, 18*R**-**1**

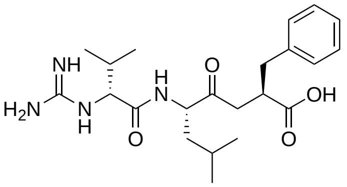

2*R**, 12*S**, 18*R**-**1**

| | C position | 2*R**, 12*S**, 18*S**-1 | 2*R**, 12*R**, 18*S**-1 | 2*R**, 12*R**, 18*R**-1 | 2*R**, 12*S**, 18*R**-1 | ketomemicin C-418 (1) |
|---|---|---|---|---|---|---|
| **$^{13}$C chemical shifts** | **1** | 184.0 | 185.9 | 183.3 | 186.5 | 182.0 |
| | **2** | 52.8 | 54.4 | 51.4 | 55.1 | 46.4 |
| | **3** | 41.0 | 39.8 | 38.1 | 36.7 | 39.6 |
| | **4** | 141.8 | 143.0 | 144.1 | 143.6 | 141.5 |
| | **5** | 130.3 | 130.1 | 130.0 | 131.0 | 130.0 |
| | **6** | 128.6 | 128.6 | 128.5 | 128.2 | 129.1 |
| | **7** | 126.8 | 126.3 | 126.0 | 126.0 | 126.9 |
| | **8** | 129.0 | 128.4 | 128.3 | 128.5 | 129.1 |
| | **9** | 129.7 | 130.4 | 130.6 | 130.0 | 130.0 |
| | **10** | 40.1 | 44.1 | 41.1 | 39.8 | 41.7 |
| | **11** | 215.8 | 215.2 | 215.7 | 215.8 | 209.4 |
| | **12** | 60.65 | 58.62 | 60.10 | 59.7 | 58.5 |
| | **13** | 41.51 | 38.17 | 41.45 | 37.8 | 39.7 |
| | **14** | 28.42 | 28.54 | 28.16 | 27.6 | 26.0 |
| | **15** | 22.90 | 23.24 | 21.12 | 21.4 | 23.6 |
| | **16** | 18.92 | 20.51 | 21.11 | 22.7 | 21.4 |
| | **17** | 169.40 | 169.74 | 170.28 | 169.8 | 171.7 |
| | **18** | 67.49 | 64.33 | 67.62 | 64.5 | 62.0 |
| | **19** | 35.05 | 32.92 | 35.21 | 33.0 | 32.1 |
| | **20** | 18.64 | 19.07 | 18.80 | 18.9 | 19.4 |
| | **21** | 17.94 | 16.25 | 18.17 | 16.5 | 17.8 |
| | **22** | 154.05 | 156.48 | 153.95 | 156.5 | 158.9 |

**Table S4.** Calculated $^{13}$C NMR chemical shifts for diastereomers of **1**. Experimental values for natural product added for reference.

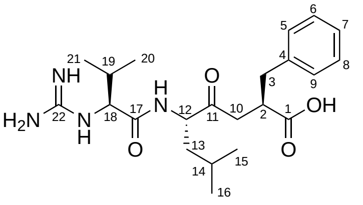

2*R**, 12*S**, 18*S**-**1**

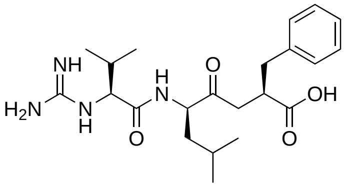

2*R**, 12*R**, 18*S**-**1**

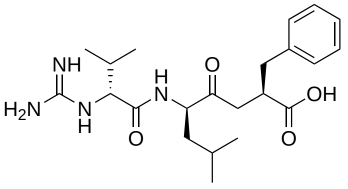

2*R**, 12*R**, 18*R**-**1**

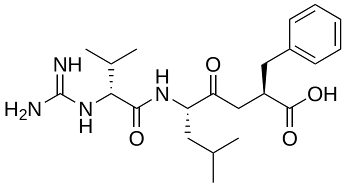

2*R**, 12*S**, 18*R**-**1**

| | **C position** | **2*R**, 12*S**, 18*S**-1** | **2*R**, 12*R**, 18*S**-1** | **2*R**, 12*R**, 18*R**-1** | **2*R**, 12*S**, 18*R**-1** | **ketomemicin C-418 (1)** |
|---|---|---|---|---|---|---|
| ***$^1$H chemical shifts*** | **2** | 2.80 | 2.61 | 2.60 | 2.52 | 3.01 |
| | **3a** | 3.04 | 3.00 | 3.09 | 2.89 | 3.05 |
| | **3b** | 2.62 | 2.52 | 2.89 | 2.80 | 2.65 |
| | **5** | 7.37 | 7.37 | 7.35 | 7.33 | 7.22 |
| | **6** | 7.39 | 7.34 | 7.32 | 7.30 | 7.25 |
| | **7** | 7.29 | 7.24 | 7.22 | 7.22 | 7.16 |
| | **8** | 7.32 | 7.31 | 7.30 | 7.32 | 7.25 |
| | **9** | 7.24 | 7.28 | 7.56 | 7.75 | 7.22 |
| | **10a** | 3.08 | 2.76 | 2.87 | 2.09 | 2.88 |
| | **10b** | 1.75 | 2.02 | 2.38 | 2.95 | 2.32 |
| | **12** | 4.21 | 4.53 | 4.29 | 4.51 | 4.43 |
| | **13a** | 1.81 | 1.78 | 1.54 | 1.69 | 1.67 |
| | **13b** | 1.42 | 1.01 | 1.48 | 1.49 | 1.47 |
| | **14** | 1.20 | 1.61 | 1.56 | 1.53 | 1.63 |
| | **15** | 0.92 | 0.88 | 0.92 | 0.94 | 0.94 |
| | **16** | 0.83 | 0.86 | 0.92 | 0.98 | 0.92 |
| | **18** | 3.16 | 3.79 | 3.13 | 3.82 | 3.88 |
| | **19** | 1.95 | 2.74 | 1.90 | 2.64 | 2.18 |
| | **20** | 1.07 | 1.10 | 1.05 | 1.08 | 0.98 |
| | **21** | 1.01 | 1.02 | 0.98 | 0.94 | 0.95 |

**Table S5.** Calculated $^1$H NMR chemical shifts for diastereomers of **1**. Experimental values for natural product added for reference.

| | *2R**, *12S**, *18S**-1 | *2R**, *12R**, *18S**-1 | *2R**, *12R**, *18R**-1 | *2R**, *12S**, *18R**-1 |
|---|---|---|---|---|
| ***$^{13}$C-MAE*** | 2.1 | 1.8 | 2.0 | 2.2 |
| ***$^{1}$H-MAE*** | 0.18 | 0.16 | 0.15 | 0.20 |
| ***overall-MAE*** | 5.6 | 4.9 | 5.0 | 6.2 |
| ***$^{13}$C -RMSD*** | 2.9 | 2.6 | 2.7 | 2.9 |
| ***$^{1}$H -RMSD*** | 0.26 | 0.21 | 0.23 | 0.30 |
| ***overall-RMSD*** | 8.0 | 6.9 | 7.3 | 9.0 |

**Table S6.** Mean averaged error (MAE) and root-mean-squared deviation (RMSD) between calculated and experimental NMR chemical shifts of **1**. The overall-MAE = $^{13}$C-MAE + 20×($^{1}$H-MAE). The overall-RMSD = $^{13}$C-RMSD + 20×($^{1}$H-RSMD).

| | | **2*R**, 12*S**, 18*S**-1** | **2*R**, 12*R**, 18*S**-1** | **2*R**, 12*R**, 18*R**-1** | **2*R**, 12*S**, 18*R**-1** |
|---|---|---|---|---|---|
| **DP4+ (%)** | **H data** | 0.73% | 11.77% | 87.12% | 0.38% |
| | **C data** | 0.36% | 98.50% | 0.91% | 0.22% |
| | **All data** | **0.02%** | **93.55%** | **6.42%** | **0.01%** |

**Table S7.** Summary of DP4+ results between calculated and experimental NMR chemical shifts of **1**.

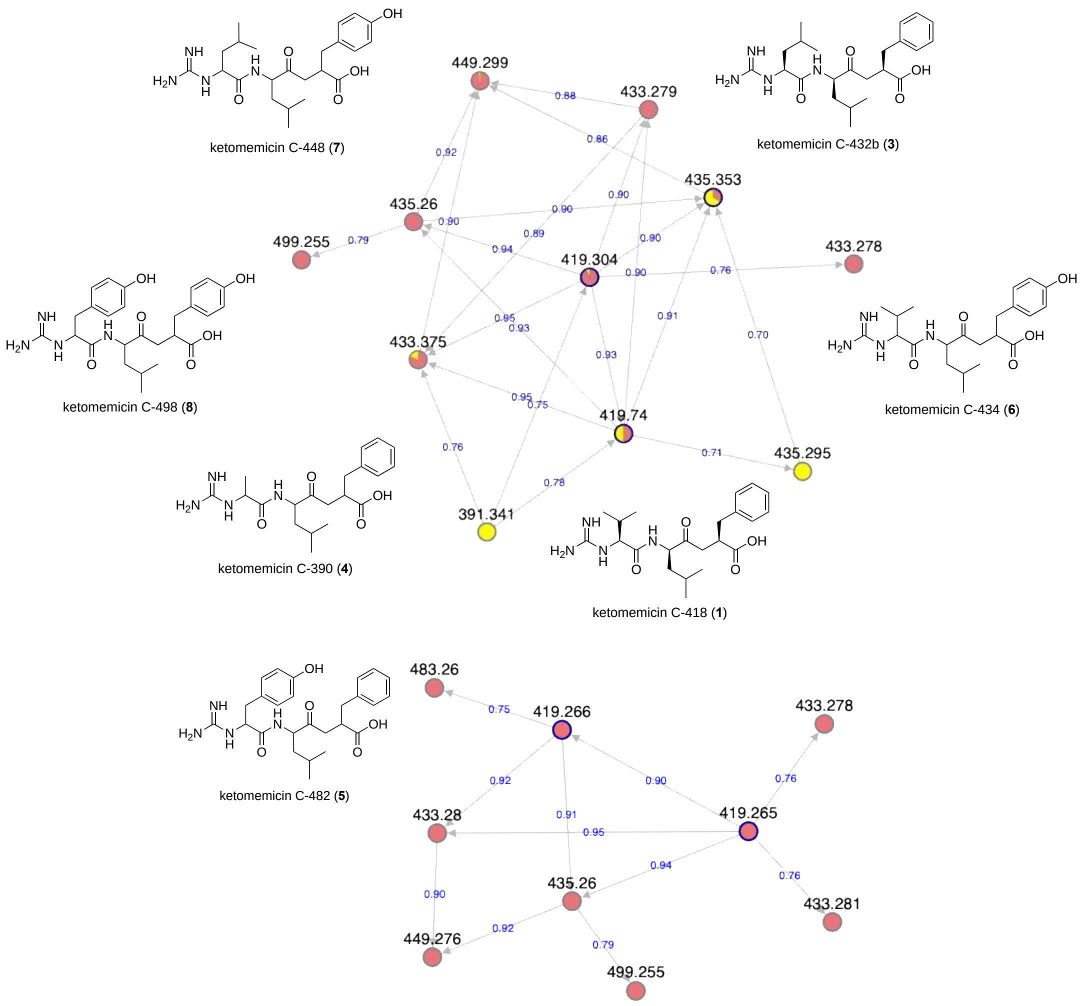


**Figure S16**. Ketomemicin-specific clusters observed with GNPS molecular networking.

GNPS “classical” analysis of an organic extract of *S. pacifica* CNY498 (yellow nodes) and the Crüsemann et al. (2017) Massive LCMS dataset (MSV000078836, red nodes). Using the same GNPS parameters, the cluster on the bottom was obtained by running MSV00078836 by itself, without the CNY498 extract LCMS data. Ketomemicin C-482 (**5**) was only detected in this latter analysis. Lines (“edges”) connecting nodes show the similarity score (in blue) between the fragmentation spectra of the corresponding nodes. The structures and names of the ketomemicin analogs identified through this analysis are shown. The node labels represent the mass of precursor $[M+H]^+$ ions.

GNPS molecular networks were ran with standard parameters (precursor ion mass tolerance: 2.0 Da, fragment ion mass tolerance: 0.5 Da, minimum matched fragment ions: 6, minimum cluster size: 2, maximum shift: 1999 Da, and minimum pair cosine: 0.70). The Duncan et. al (2016) (MSV000079284 and MSV000079484) and Chase et al. (2021) (MSV000085890) Massive datasets were also analyzed but these showed low or nil presence of ketomemicins.

| | | | ketomemicin C-418 (**1**) | | | ketomemicin C-432 (**2** - **3**) | | | ketomemicin C-482 (**5**) | | | ketomemicin C-334 (**6**) | | | ketomemicin C-348 (**7**) | | | ketomemicin C-498 (**8**) | | | |
|---|---|---|---|---|---|---|---|---|---|---|---|---|---|---|---|---|---|---|---|---|---|
| | | | **419.2653 *m/z*** | | | **433.2809 *m/z*** | | | **483.2602 *m/z*** | | | **435.2602 *m/z*** | | | **449.2758 *m/z*** | | | **499.2551 *m/z*** | | | |
| | | | **intensity** | **$t_R$** | **MS/MS** | **intensity** | **$t_R$** | **MS/MS** | **intensity** | **$t_R$** | **MS/MS** | **intensity** | **$t_R$** | **MS/MS** | **intensity** | **$t_R$** | **MS/MS** | **intensity** | **$t_R$** | **MS/MS** | |
| | **118 Salinispora strains** | | | | | | | | | | | | | | | | | | | | **ktm BGC** |
| 1 | CNY012 | tropica | 2.1E+03 | 12.62 | 142.096 | | | | | | | 5.3E+04 | 11.34 | 142.098 | 2.2E+04 | 11.91 | 156.112 | 6.7E+03 | 11.37 | 206.09 | yes |
| 2 | CNH898 | tropica | | | | | | | | | | 2.3E+03 | 11.47 | 142.097 | | | | | | | yes |
| 3 | CNB536 | tropica | | | | | | | | | | | | | | | | | | | yes |
| 4 | CNT261 | tropica | | | | | | | | | | | | | | | | | | | yes |
| 5 | CNB476 | tropica | | | | | | | | | | 3.0E+04 | 11.35 | 142.095 | 1.5E+04 | 11.79 | 156.11 | 7.1E+03 | 11.41 | 206.09 | yes |
| 6 | CNS416 | tropica | | | | | | | | | | | | | | | | | | | yes |
| 7 | CNT250 | tropica | 2.1E+03 | 12.9 | 142.096 | | | | | | | 2.3E+04 | 11.56 | 142.097 | 7.1E+03 | 12.01 | 156.112 | | | | yes |
| 8 | CNS197 | tropica | | | | | | | | | | 2.5E+04 | 11.27 | 142.095 | 1.1E+04 | 11.85 | 156.112 | 2.6E+03 | 11.34 | 206.089 | yes |
| 9 | CNR699 | tropica | | | | | | | | | | 1.1E+04 | 11.37 | 142.094 | 4.0E+03 | 11.91 | 156.108 | | | | yes |
| 10 | CNB440 | tropica | | | | | | | | | | 3.6E+04 | 11.41 | 142.095 | 8.4E+03 | 11.86 | 156.113 | | | | yes |
| 11 | CNY681 | tropica | | | | | | | | | | | | | | | | | | | yes |
| 12 | CNY678 | tropica | | | | | | | | | | 3.7E+03 | 11.36 | 142.095 | 3.6E+03 | 11.91 | 156.112 | | | | yes |
| 1 | CNR942 | fenicalii | | | | | | | | | | | | | | | | | | | yes |
| 2 | CNT569 | fenicalii | | | | | | | 7.9E+03 | 12.75 | 206.09 | | | | | | | 9.4E+03 | 12.23 | 206.093 | yes |
| 1 | CNY202 | cortesiana | | | | | | | | | | | | | | | | | | | yes |
| 1 | CNY646 | mooreana | | | | | | | | | | | | | | | | | | | yes |
| 2 | CNS237 | mooreana | 1.1E+05 | 12.79 | 142.096 | 6.9E+04 | 13.24 | 156.112 | | | | | | | | | | | | | yes |
| 3 | CNT150 | mooreana | | | | | | | | | | | | | | | | | | | partial |
| 1 | CNT854 | oceanensis | | | | | | | | | | | | | | | | | | | yes |
| 2 | CNT584 | oceanensis | | | | 5.9E+03 | 12.77 | 156.111 | | | | | | | | | | | | | yes |
| 3 | CNT124 | oceanensis | | | | | | | | | | | | | | | | | | | yes |
| 4 | CNY673 | oceanensis | | | | | | | | | | | | | | | | | | | no |
| 5 | CNY703 | oceanensis | | | | | | | | | | | | | | | | | | | no |
| 6 | CNT138 | oceanensis | | | | | | | | | | | | | | | | | | | no |
| 7 | CNT029 | oceanensis | | | | | | | | | | | | | | | | | | | no |
| 8 | CNT045 | oceanensis | | | | | | | | | | | | | | | | | | | no |
| 9 | CNS996 | oceanensis | | | | | | | | | | | | | | | | | | | no |
| 10 | CNT403 | oceanensis | | | | | | | | | | | | | | | | | | | no |
| 11 | CNS863 | oceanensis | | | | | | | | | | | | | | | | | | | no |
| 12 | CNS860 | oceanensis | | | | | | | | | | | | | | | | | | | no |
| 1 | CNY666 | goodfellowii | | | | | | | | | | | | | | | | | | | yes |
| 1 | CNS055 | vitiensis | | | | | | | | | | | | | | | | | | | no |
| 2 | CNT148 | vitiensis | | | | | | | | | | | | | | | | | | | no |
| 3 | CNS801 | vitiensis | | | | | | | | | | | | | | | | | | | no |
| 1 | CNT609 | pacifica | | | | | | | | | | | | | | | | | | | yes |
| 2 | CNT133 | pacifica | | | | | | | | | | | | | | | | | | | yes |
| 3 | CNT084 | pacifica | | | | | | | | | | | | | | | | | | | yes |
| 4 | CNT851 | pacifica | | | | | | | | | | | | | | | | | | | partial |
| 5 | CNT796 | pacifica | | | | | | | | | | | | | | | | | | | partial |
| 6 | CNY330 | pacifica | | | | | | | | | | | | | | | | | | | yes |
| 7 | CNY331 | pacifica | | | | | | | | | | | | | | | | | | | yes |
| 8 | CNY363 | pacifica | | | | | | | | | | | | | | | | | | | yes |
| 9 | CNT855 | pacifica | | | | | | | | | | | | | | | | | | | yes |
| 10 | CNY498 | pacifica | | | | 2.6E+03 | 13.14 | 156.11 | | | | | | | | | | | | | yes |
| 11 | CNS960 | pacifica | 2.7E+03 | 12.89 | 142.097 | 4.3E+03 | 13.34 | 156.11 | | | | | | | | | | | | | yes |
| 12 | CNT003 | pacifica | | | | 2.3E+03 | 14.02 | 156.115 | | | | | | | | | | | | | yes |
| 13 | CNR114 | pacifica | | | | 9.7E+03 | 12.7 | 156.112 | | | | | | | | | | | | | yes |
| 14 | CNQ768 | pacifica | 3.9E+03 | 12.9 | 142.098 | 4.5E+03 | 13.38 | 156.112 | | | | | | | | | | | | | yes |
| 15 | CNS103 | pacifica | | | | | | | | | | | | | | | | | | | yes |
| 16 | CNR909 | pacifica | 4.8E+03 | 13.15 | 142.096 | 3.9E+03 | 13.64 | 156.112 | | | | | | | | | | | | | yes |
| 17 | CNH732 | pacifica | | | | | | | | | | | | | | | | | | | yes |
| 18 | CNR510 | pacifica | 2.5E+03 | 13.02 | 142.098 | 2.6E+03 | 13.49 | 156.11 | | | | | | | | | | | | | yes |
| 19 | CNR894 | pacifica | 1.1E+04 | 12.8 | 142.098 | 1.3E+04 | 13.23 | 156.112 | | | | | | | | | | | | | yes |
| 20 | CNT131 | pacifica | 1.5E+04 | 12.46 | 142.096 | 1.5E+04 | 12.93 | 156.113 | | | | | | | | | | | | | yes |
| 21 | CNT603 | pacifica | 2.3E+04 | 12.6 | 142.098 | 1.9E+04 | 12.98 | 156.112 | | | | | | | | | | | | | yes |
| 22 | CNT001 | pacifica | 6.3E+03 | 12.46 | 142.097 | 4.6E+03 | 12.89 | 156.11 | | | | | | | | | | | | | yes |
| 23 | CNY239 | pacifica | | | | | | | | | | | | | | | | | | | yes |

**Table S8.** Distribution of ketomemicin analogs (**1–8**) and ketomemicin biosynthetic gene cluster (*ktm*) in 118 *Salinispora* strains.

| | | | ketomemicin C-418 (**1**) | | | ketomemicin C-432 (**2** - **3**) | | | ketomemicin C-482 (**5**) | | | ketomemicin C-334 (**6**) | | | ketomemicin C-348 (**7**) | | | ketomemicin C-498 (**8**) | | | |
|---|---|---|---|---|---|---|---|---|---|---|---|---|---|---|---|---|---|---|---|---|---|
| | | | **419.2653 *m/z*** | | | **433.2809 *m/z*** | | | **483.2602 *m/z*** | | | **435.2602 *m/z*** | | | **449.2758 *m/z*** | | | **499.2551 *m/z*** | | | |
| | | | **intensity** | **$t_R$** | **MS/MS** | **intensity** | **$t_R$** | **MS/MS** | **intensity** | **$t_R$** | **MS/MS** | **intensity** | **$t_R$** | **MS/MS** | **intensity** | **$t_R$** | **MS/MS** | **intensity** | **$t_R$** | **MS/MS** | |
| | **118 Salinispora strains** | | | | | | | | | | | | | | | | | | | | **ktm BGC** |
| 1 | CNB527 | arenicola | | | | | | | | | | | | | | | | | | | partial |
| 2 | CNY685 | arenicola | | | | | | | | | | | | | | | | | | | partial |
| 3 | CNY690 | arenicola | | | | | | | | | | | | | | | | | | | partial |
| 4 | CNY694 | arenicola | | | | | | | | | | | | | | | | | | | partial |
| 5 | CNB458 | arenicola | | | | | | | | | | | | | | | | | | | partial |
| 6 | CNH962 | arenicola | | | | | | | | | | | | | | | | | | | partial |
| 7 | CNH963 | arenicola | | | | | | | | | | | | | | | | | | | partial |
| 8 | CNY486 | arenicola | | | | | | | | | | | | | | | | | | | partial |
| 9 | CNH996B | arenicola | | | | | | | | | | | | | | | | | | | partial |
| 10 | CNH996 | arenicola | | | | | | | | | | | | | | | | | | | partial |
| 11 | CNH713 | arenicola | | | | | | | | | | | | | | | | | | | partial |
| 12 | CNH941 | arenicola | | | | | | | | | | | | | | | | | | | no |
| 13 | CNP193 | arenicola | | | | | | | | | | | | | | | | | | | no |
| 14 | CNH964 | arenicola | | | | | | | | | | | | | | | | | | | no |
| 15 | CNP105 | arenicola | | | | | | | | | | | | | | | | | | | no |
| 16 | CNY011 | arenicola | | | | | | | | | | | | | | | | | | | partial |
| 17 | CNH877 | arenicola | | | | | | | | | | | | | | | | | | | partial |
| 18 | CNY679 | arenicola | | | | | | | | | | | | | | | | | | | partial |
| 19 | CNH905 | arenicola | | | | | | | | | | | | | | | | | | | partial |
| 20 | CNH646 | arenicola | | | | | | | | | | | | | | | | | | | partial |
| 21 | CNH643 | arenicola | | | | | | | | | | | | | | | | | | | partial |
| 22 | CNS991 | arenicola | | | | | | | | | | | | | | | | | | | yes |
| 23 | CNS848 | arenicola | | | | | | | | | | | | | | | | | | | yes |
| 24 | CNY280 | arenicola | 2.3E+03 | 12.64 | 142.092 | 3.0E+03 | 13.12 | 156.113 | | | | | | | | | | | | | yes |
| 25 | CNT857 | arenicola | | | | | | | | | | | | | | | | | | | yes |
| 26 | CNT859 | arenicola | | | | | | | | | | | | | | | | | | | yes |
| 27 | CNT850 | arenicola | | | | | | | | | | | | | | | | | | | yes |
| 28 | CNT799 | arenicola | | | | | | | | | | | | | | | | | | | yes |
| 29 | CNT849 | arenicola | | | | | | | | | | | | | | | | | | | partial |
| 30 | CNT798 | arenicola | | | | | | | | | | | | | | | | | | | partial |
| 31 | CNT800 | arenicola | | | | | | | | | | | | | | | | | | | partial |
| 32 | CNX482 | arenicola | | | | | | | | | | | | | | | | | | | yes |
| 33 | CNX481 | arenicola | | | | | | | | | | | | | | | | | | | yes |
| 34 | CNX891 | arenicola | | | | 2.7E+03 | 13.3 | 156.114 | | | | | | | | | | | | | yes |
| 35 | CNX508 | arenicola | | | | | | | | | | | | | | | | | | | yes |
| 36 | CNX814 | arenicola | | | | | | | | | | | | | | | | | | | yes |
| 37 | CNH718 | arenicola | | | | | | | | | | | | | | | | | | | yes |
| 38 | CNS299 | arenicola | | | | | | | | | | | | | | | | | | | yes |
| 39 | CNR921 | arenicola | | | | | | | | | | | | | | | | | | | yes |
| 40 | CNS243 | arenicola | | | | | | | | | | | | | | | | | | | yes |
| 41 | CNR107 | arenicola | | | | 2.1E+03 | 13.16 | 156.111 | | | | | | | | | | | | | yes |
| 42 | CNQ884 | arenicola | | | | | | | | | | | | | | | | | | | yes |
| 43 | CNS325 | arenicola | | | | | | | | | | | | | | | | | | | yes |
| 44 | CNS205 | arenicola | | | | | | | | | | | | | | | | | | | yes |
| 45 | CNS296 | arenicola | | | | | | | | | | | | | | | | | | | yes |
| 46 | CNS051 | arenicola | | | | | | | | | | | | | | | | | | | yes |
| 47 | CNR425 | arenicola | | | | | | | | | | | | | | | | | | | yes |
| 48 | CNQ748 | arenicola | | | | | | | | | | | | | | | | | | | yes |
| 49 | CNT005 | arenicola | | | | | | | | | | | | | | | | | | | yes |
| 50 | CNS744 | arenicola | | | | | | | | | | | | | | | | | | | yes |
| 51 | CNY231 | arenicola | | | | | | | | | | | | | | | | | | | yes |
| 52 | CNY256 | arenicola | | | | | | | | | | | | | | | | | | | yes |
| 53 | CNS673 | arenicola | | | | | | | | | | | | | | | | | | | yes |
| 54 | CNY260 | arenicola | | | | | | | | | | | | | | | | | | | yes |
| 55 | CNS820 | arenicola | | | | | | | | | | | | | | | | | | | yes |
| 56 | CNY237 | arenicola | | | | | | | | | | | | | | | | | | | yes |
| 57 | CNY230 | arenicola | | | | | | | | | | | | | | | | | | | yes |
| 58 | CNY244 | arenicola | | | | | | | | | | | | | | | | | | | yes |
| 59 | CNS342 | arenicola | | | | | | | | | | | | | | | | | | | yes |
| 60 | CNY234 | arenicola | | | | | | | | | | | | | | | | | | | yes |
| 61 | CNY282 | arenicola | | | | | | | | | | | | | | | | | | | yes |

**Table S8 (continued).** Distribution of ketomemicin analogs (**1–8**) and ketomemicin biosynthetic gene cluster (*ktm*) in 118 *Salinispora* strains.

Masses of ketomemicin analogs were searched in the Crüsemann et al. (2017) LCMS dataset using MZmine to assess distribution of these metabolites in 118 *Salinispora* strains. Data on the extracted masses, including peak intensity values (color coded to represent the production levels: high (green, $1x10^5$ counts), low (red, $2x10^3$ counts) or intermediate (light green, yellow, or orange for decreasing values)), retention times ($t_R$, noted in minutes), and characteristic MS/MS fragment ions, are shown. Ketomemicin C-390 (**4**) was only detected in the extract of *S. pacifica* CNY-498 from which **1–3** were isolated. Strains are in phylogenomic order based on the results of Román-Ponce et al. (2020).

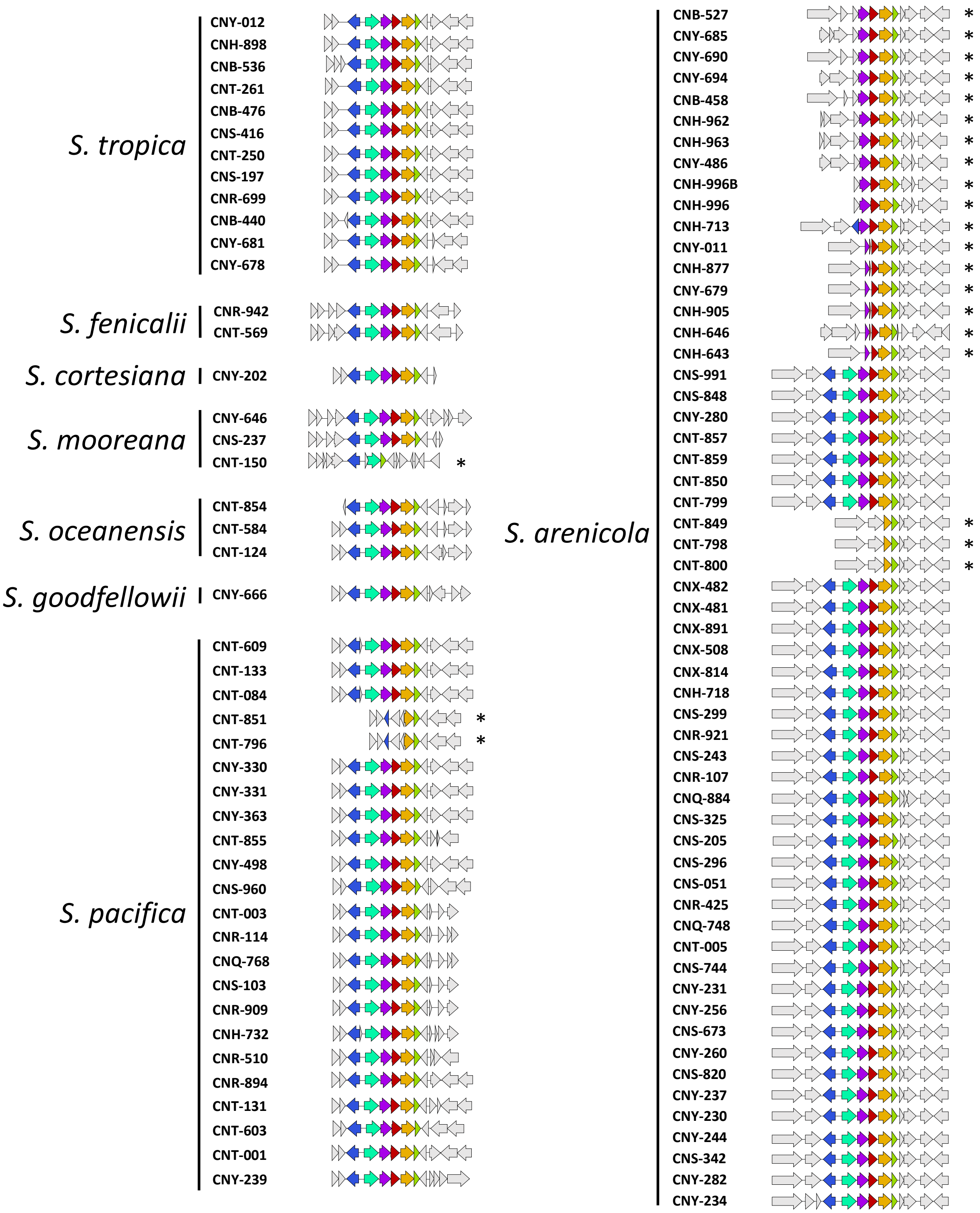


**Figure S17.** Ketomemicin biosynthetic gene clusters (*ktm*) in *Salinispora* genomes.

*Ktm* homologs identified using Cblaster were then visualized using Clinker. *Ktm* BGCs were identified in 103 of 118 *Salinispora* genomes. Genes are colored as: *ktmA* (red), *ktmB* (yellow), *ktmC* (olive), *ktmD* (cyan), *ktmE* (blue), and *ktmF* (purple). Asterisks (*) denote partial clusters with two to five *ktm* genes (23 BGCs). Strains are in phylogenomic order based on the results of Román-Ponce et al. (2020).

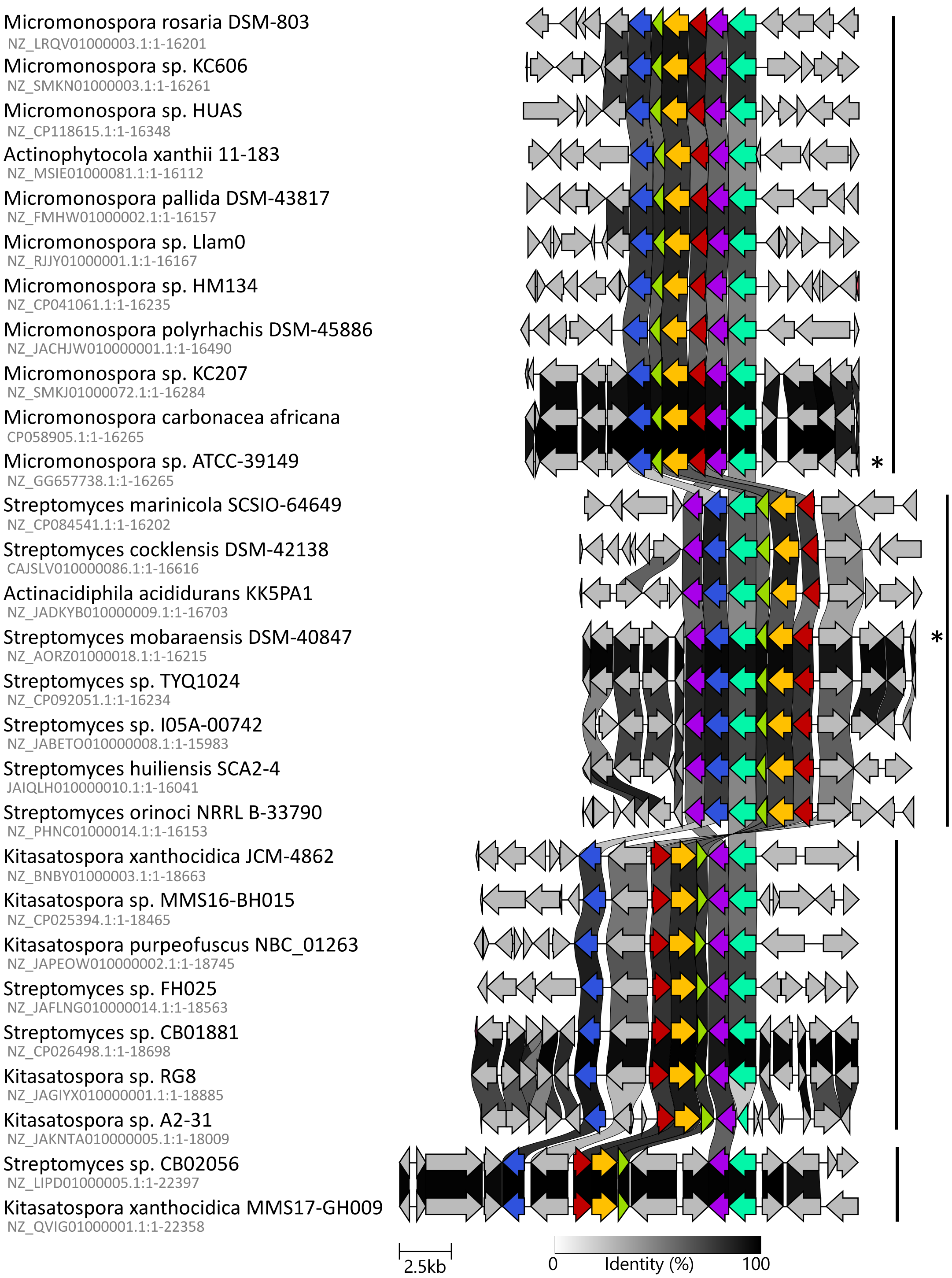


**Figure S18.** Synteny plot of ketomemicin-like biosynthetic gene clusters (*ktm*) in the NCBI database (excluding *Salinispora*)

BGCs were identified using Cblaster (with parameters: 30% identity between gene homologs and 30% coverage, minimum 6 matching genes) by searching for *ktmA*–*ktmF* homologs from *S. pacifica* CNY-498 against the NCBI RefSeq and Non-redundant databases. Hits shared by the two databases and hits within *Salinispora* were manually removed. *Ktm* homologs are color coded: *ktmA* (red), *ktmB* (yellow), *ktmC* (olive), *ktmD* (cyan), *ktmE* (blue), and *ktmF* (purple). BGCs with biochemical validation are highlighted with an asterisk (*). Vertical bars denote the four types of *ktm*-like BGCs observed.
